# Whole-genome DNA methylation profiling in COVID-19 positive patients reveals alterations in pathways linked to neurological dysfunction

**DOI:** 10.1101/2025.09.25.678349

**Authors:** Saima Zameer, Ehraz Anis, Qiang Sha, Martha L. Escobar Galvis, Sanam Khan, Jennifer A. Steiner, Milda Milčiūtė, Ieva Kerševičiūtė, Migle Gabrielaite, Juozas Gordevicius, Andrew Pospisilik, Nazia Saiyed, Stewart F. Graham, Patrik Brundin, Lena Brundin

**Author notes:** Corresponding author: Lena Brundin Address: Department of Neurodegenerative Science, Van Andel Institute 333 Bostwick Ave NE, Grand Rapids, MI 49503, USA.

## Abstract

**Background:** Severe acute respiratory syndrome coronavirus 2 (SARS-CoV-2) is a highly transmissible RNA betacoronavirus, causing coronavirus disease-19 (COVID-19). Infection with SARS-CoV-2 can result in a broad spectrum of clinical outcomes, ranging from asymptomatic or mild to a severe, deadly illness. Emerging evidence suggests SARS-CoV-2 affects host gene regulation through epigenetic mechanisms, such as DNA methylation, potentially contributing to immune dysregulation and post-acute sequelae, including neurological and psychiatric disorders. However, the extent and functional relevance of these epigenetic changes remain uncertain.

**Methods and results:** We employed whole-genome bisulfite sequencing to profile DNA methylation in peripheral blood from SARS-CoV-2-positive patients across a spectrum of symptom severity, ranging from asymptomatic to severe (n=101), in comparison to SARS-CoV-2-negative individuals (n=105). We observed a widespread hypomethylation in the genomes of infected individuals, which was more pronounced in severe cases. Notably, we identified differentially methylated genes in patients with mild (19 genes), moderate (19 genes), and severe (35 genes) symptoms. These genes included those involved in canonical immune responses as well as known to be linked to neurodegenerative diseases. Subsequent pathway enrichment analysis further supported the significant association between the differentially methylated genes and those implicated in Alzheimer’s and Parkinson’s disease, as well as neuropsychiatric conditions, suggesting potential epigenetic links between acute SARS-CoV-2 infection and long-term neurological outcomes. This is one of the first studies to comprehensively map severity-stratified genome-wide DNA methylation changes in COVID-19 patients.

**Conclusion:** Our findings underscore the potential importance of epigenetic regulation in the acute responses to SARS-CoV-2 infection and highlight an overlap with epigenetic mechanisms relevant for neuropsychiatric disease processes.

## Introduction

Over 7 million people have died worldwide due to the severe acute respiratory syndrome coronavirus 2 (SARS-CoV-2; species: *Betacoronavirus pandemicum*) as of 2025 (1, 2). This novel RNA betacoronavirus, first detected in humans in the fall of 2019, led to a highly transmissible illness called coronavirus disease 19 (COVID-19) that quickly developed into a global pandemic (3, 4). COVID-19 exhibits a wide range of clinical symptoms, from asymptomatic or mild cases to severe, life-threatening systemic illness (5, 6), It presents with significant variation in severity and symptoms among patients. Common symptoms include cough, fever, chest tightness, shortness of breath, anosmia, and ageusia. Many cases also exhibit gastrointestinal, neurological, and cardiovascular symptoms (7, 8). The clinical outcomes of COVID-19, including the risk of death, are partly related to comorbidities such as hypertension, diabetes, and cardiovascular disease, as well as demographic factors, especially age, sex, and ethnicity. The highest risk of death is associated with the development of interstitial pneumonia, dyspnea, and acute respiratory distress syndrome (9, 10).

While understanding virus-host interactions is crucial for disease dynamics, variations in the SARS-CoV-2 genome can also help explain the wide range of clinical outcomes seen. However, new evidence indicates that differences in human biology, rather than viral traits, have a greater impact on disease severity. For example, even among different viral strains, the variation in how hosts respond is much greater than the differences caused by viral genetic diversity (11, 12).

The significant variability among individuals in the severity of COVID-19 has created major challenges in understanding its biological foundations (13) Genome-wide association studies (GWAS) have pinpointed specific gene loci linked to COVID-19-related phenotypes. These studies emphasize genes involved in viral entry, immune responses (both innate and adaptive), as well as pathways associated with lung function and systemic inflammation (14–17). Along with genetic predisposition, certain epigenetic changes are also recognized as important factors that influence susceptibility and severity (18, 19). Several reports suggest that SARS-CoV-2 can induce significant epigenetic reprogramming, such as DNA methylation and histone modifications, which influence immune cell function and inflammatory responses (4, 20, 21). Besides impacting the individual’s immune system, SARS-CoV-2 has been shown to directly infect non-immune and non-respiratory tissues and cells, further complicating the disease’s systemic effects and potential long-term consequences (22).

Despite these insights, the relationship between host epigenetic changes and the clinical severity of COVID-19 remains poorly understood. A better understanding of epigenetic alterations could provide novel biomarkers for disease prognosis and reveal mechanisms linking viral infection to long-term complications, including neurological outcomes. Therefore, this study aimed to investigate genome-wide DNA methylation patterns in SARS-CoV-2-positive patients compared to SARS-CoV-2-negative controls, to identify epigenetic signatures associated with disease severity.

## Materials and Methods

### Clinical Cohort

This study was approved by the Institutional Review Boards at Corewell Health East William Beaumont University Hospital (formerly Beaumont Health), Royal Oak, MI, USA (approval no. 2020-269) and Van Andel Institute, Grand Rapids, MI, USA (approval no. NHS 21014). The SARS-CoV-2-positive cohort was divided into three groups based on hospital admission status and the maximum level of oxygen administered during admission: 1) Asymptomatic or Mild – non-hospitalized patients with no oxygen required, 2) Moderate – hospitalized patients receiving nasal cannula at ≤6L of oxygen, and 3) Severe – hospitalized patients using nasal cannula >6L, or on O2 devices such as ventilation masks, cold high-flow, heated high-flow, non-rebreather masks, Bi-PAP, or CPAP. These hospital-based controls included Beaumont Health staff or patients who tested negative for SARS-CoV-2. SARS-CoV-2-positive patients were defined as those who tested positive via routine Polymerase Chain Reaction (PCR) testing of a nasopharyngeal swab, while controls tested negative.

### Human Samples

Blood samples were collected from 101 SARS-CoV-2-positive patients (44 with asymptomatic to mild symptoms, 37 with moderate symptoms, and 20 with severe symptoms), as well as 105 hospital-based SARS-CoV-2-negative controls, during 2021. The patients’ comorbidities were documented through their clinical history. Samples were collected in EDTA tubes and immediately stored at 4°C, then at -80°C until analysis. The initial cohort included 206 samples, with seven samples removed due to poor quality (Figure 1).

**Figure 1.**
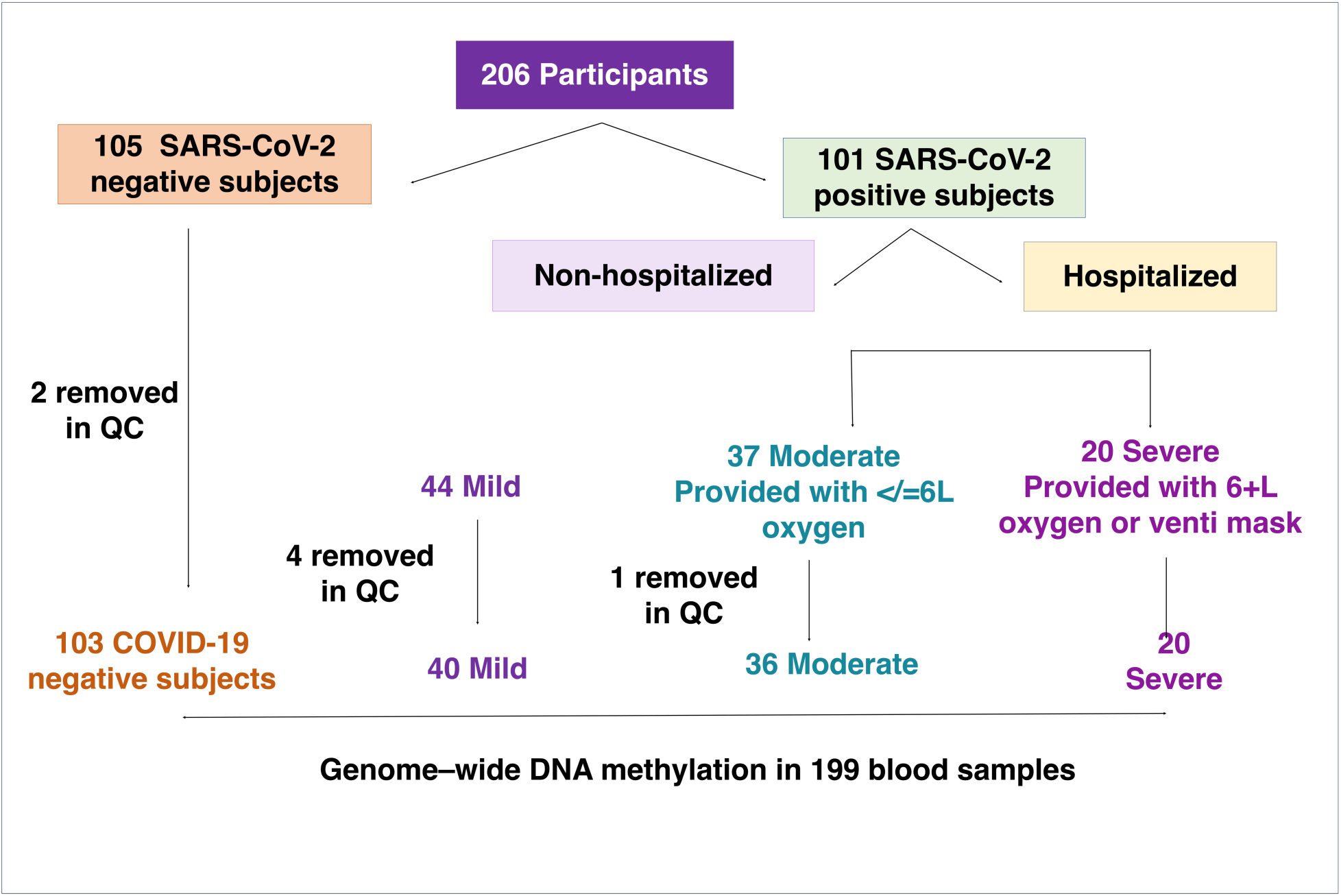
SARS-CoV-2-positive and SARS-CoV-2-negative cohorts. A total of 206 participants were initially enrolled; however, 11 individuals were excluded from the analysis due to inadequate data quality or missing information. QC - Quality Control and WGBS – Whole-Genome Bisulfite Sequencing.

### DNA extraction and whole genome sequence libraries generation

DNA from blood samples was extracted using the QiAamp DNA Blood Mini kit (Qiagen Cat No. #51106) following the manufacturer’s protocol. DNA concentration was measured with a Nanodrop spectrophotometer (ThermoFisher). Next, the extracted DNA samples were prepared for whole-genome sequencing library construction. DNA controls, pUC19 and unmethylated lambda DNA (0.0005% and 0.01%, respectively), were added to each high-molecular-weight genomic DNA sample. All prepared libraries involved shearing the DNA to an average size of approximately 350 bp. Libraries were prepared from the NEBNext Enzymatic Methyl-seq Kit (New England Biolabs, Cat. #E7120L) using 200 ng of sheared DNA as input, and libraries were constructed following the kit protocol. The denaturation was performed using 0.1 N sodium hydroxide, followed by four cycles of PCR amplification. KAPA pure beads (Roche Sequencing, Cat. #KK8001) were used for cleanup steps in all prepared libraries. Quality and quantity of the finished libraries were evaluated using a combination of Agilent High Sensitivity DNA chip (Agilent Technologies, Inc., Cat. #5067-4626), QuantiFluor® dsDNA System (Promega Corp., Cat. #E2670), and Kapa Illumina Library Quantification qPCR assay (Roche Sequencing, Cat. # KK4824). 150 bp paired-end sequencing was performed on an Illumina NovaSeq 6000 sequencer using an S4, 300 bp sequencing kit (Illumina Inc., San Diego, CA, USA), with 10% PhiX. Illumina RTA3 did base calling, and the output of NCS was demultiplexed and converted to FASTQ format with Illumina Bcl2fastq v1.9.0.

### Data Processing and Quality Control

The raw sequencing data were preprocessed using fastp (v0.23.2) (23) for quality control. Adapter sequences were trimmed after auto-detection, along with the first 5 bases from both ends of the reads. Reads were retained if they had an average Phred quality score of ≥20 and a minimum length of 50 bp. High-quality reads were aligned to the human reference genome (GRC38, patch 13) using Arioc (v1.50) (24). Following this alignment, BAM files were processed with Sambamba (25) for read filtering (retaining reads with mapping quality ≥30), deduplication, sorting, and indexing. Per-base CpG methylation metrics were then assessed using MethylDackel (v0.6.1), and reads with low coverage were excluded (one sample was excluded due to insufficient read depth). Only CpG sites with a coverage between 15 to 500 reads were retained for downstream analysis. Coverage was calculated in 200 base pair bins across all the samples. Only regions present in at least 75% of SARS-CoV-2-positive samples (stratified by mild, moderate, and severe symptoms) were selected for the downstream methylation analysis. Beta values were computed for each tiled window for the estimation of differential methylation, which represents the degree of hypomethylation and hypermethylation relative to SARS-CoV-2-negative controls (18, 26) (Supplementary Figure S1). Missing beta values were calculated using the k-nearest neighbor imputation method with k=10 as a value for the nearest neighbors.

Post beta values calculation, pairwise sample correlations were computed, and samples with mean correlation deviating by more than two standard deviations (SDs) from the overall mean were removed as outliers (n = 3). Additionally, quality control checks were performed using MultiQC (v1.12) on both raw and preprocessed reads. To further refine the datasets, we applied Principal Component Analysis (PCA) and Multi-Dimensional Scaling (MDS) on the centered, unnormalized beta value matrix. Samples more than two SDs from the mean of the first three principal components were considered outliers and excluded from subsequent analysis (n = 3) (Supplementary Figure S2).

### Differential methylation analysis

Differentially methylated tiling windows (DMTWs) across the entire genome were initially screened in all SARS-CoV-2-positive cohorts *versus the* SARS-CoV-2-negative cohort, followed by analysis in groups stratified based on disease severity: mild, moderate, and severe *versus* negative controls. This approach allowed for the assessment of how varying levels of disease severity associate with DNA methylation patterns. Like quality control, the CpGs were divided into non-overlapping 200 bp windows, and windows with missing values in more than 50% of samples in each group were excluded from the model. Filtered tiled windows were then used to fit the linear Dispersion Shrinkage for Sequencing data (DSS) models using the R package methylSig (v1.17.0) (27). The model was adjusted for covariates including COVID-19 condition, batch, sex assigned at birth, age, race, diabetes, hypertension, cardiovascular disease, chronic obstructive pulmonary disorder (COPD), chronic kidney disease, obesity, cancer, depression, and corticosteroids. To account for unknown confounders, we used variable adjustment with the RUVg method (assuming variation is not influenced by the conditions of interest in the negative control group) from the RUVseq (v.1.36.0) R package (28), incorporating two RUVg vectors to capture unwanted variation. This adjustment was performed using the imputed beta values (a measure of methylation obtained during the quality control step), comprising high-quality windows with no missing values. Subsequently, model fitting was conducted using the diff_dss_fit function from the methylSig R package (v1.17.0).

These DMTWs were then annotated to genes using the Bioconductor package annotatr (v.1.28.0) (29) R package. When a single DMTW was associated with multiple genes, the gene symbol most frequently overlapping the region was selected for downstream analysis. The genes with DMTW are hereafter defined as differentially methylated genes (DMGs).

### Pathway enrichment analysis

To gain insight into the biological pathways and disease mechanisms affected by the differential DNA methylation patterns in SARS-CoV-2-positive patients, we computed pathway enrichment analysis using gene set enrichment analysis (GSEA) software (30) available from gsea-msigdb.org, which requires a ranked list of genes as input. The genes were ranked based on a score defined as −1×𝑠𝑖𝑔𝑛(𝐹𝐶)×𝑙𝑜𝑔(𝑝), where FC denotes differential methylation fold change and p represents the associated p-value from differential methylation analysis. If multiple methylation sites corresponding to the same gene were measured, the site with the smallest p-value was retained. We employed the human all pathway GMT file obtained from Bader Lab, Toronto, in August 2022 (31), to define gene sets for pathway enrichment analysis. This GMT file encompasses a comprehensive collection of curated gene sets derived from a wide range of databases including Gene Ontology (GO), Kyoto Encyclopedia of Gene and Genome (KEGG), Protein Analysis Through Evolutionary Relationships (PANTHER), Wiki Pathway, Reactome, Molecular Signature Database (MSigDB), NetPath, Pathbank, Encyclopedia of Human Genes and Metabolism (HumanCyc), National Cancer Institute Nature Pathway (NCL_Nature). For the enrichment score and Normalized Enrichment Score (NES), we applied the classical enrichment scoring scheme with mean-div normalization. The testing was performed for pathways with a minimum of 10 and a maximum of 500 genes. The statistical significance was assessed using 1000 permutations.

### Intersection of DMTWs, DMGs, and pathways across COVID-19 severity groups

We assessed the overlap of DMTWs, DMGs, and enriched biological pathways across mild, moderate, and severe symptom groups. To identify overlapping features among these groups, we applied statistical threshold of FDR < 0.05. Given that multiple DMTWs were mapped to a single gene, we removed redundancy by retaining only the most statistically significant DMTW per gene. Similarly, where multiple genes were associated with a single region, we assigned the gene symbol that occurred most frequently. Gene assignments and pathway enrichment analyses were performed using the rGREAT (v2.4.0) R package (32). Venn diagrams were used to visualize the intersection of DMTWs, DMGs, and pathways across different severity groups.

### Correlation analysis

To evaluate the consistency of epigenetic changes across different disease severities, we examined the correlation of DMTWs, gene order, and pathway enrichment among SARS-CoV-2-positive patients with mild, moderate, and severe symptoms. Gene order was assigned to each tiling window using the rGREAT annotation tool. Since multiple windows could be assigned to the same gene, duplicates were removed, and only the most significant DMTW for each gene was kept. Gene lists were then sorted based on the sign-logp.

### Plots generation

Manhattan plots were created using the ggplot2 R package (v3.5.2) (33) to display the genomic distribution of methylation signals, with each point representing a methylation window. The x-axis denotes genomic loci while the y-axis shows signed -log₁₀ p-values. Significant genes were annotated directly on the plots. Volcano plots were also generated using ggplot2 to depict the overall distribution and significance of methylation differences across COVID-19 severity groups. Bar plots were used to represent the proportion of hypermethylated, hypomethylated, and non-significant regions in each contrast. To explore functional implications, pathway enrichment maps were generated using the aPEAR R package (v1.0.0) (34), which clustered the top 100 GSEA-enriched pathways based on functional similarity. Clusters were labeled using the most central pathway as determined by the PageRank algorithm (35). Node color reflects the normalized enrichment score (NES) with positive values indicating a tendency for hypermethylation and negative values for hypomethylation, while node size represents the number of genes associated with each pathway and present in the input gene list. Bubble plots were used to highlight the most significantly enriched pathways related to neurodegenerative and psychiatric conditions, with bubble size corresponding to gene set size and color indicating p-value significance. Finally, heatmaps were generated using the heatmap R package (v1.0.12) to visualize correlations across SARS-CoV-2 severity groups (mild, moderate, and severe) for three features: DMTWs, gene order, and pathway enrichment.

### Statistical analysis

All data from SARS-CoV-2-positive and negative cohorts were examined for the influence of demographic and clinical covariates and were adjusted for age in all models. Initially, a generalized linear regression model was fitted using all covariates and a random number as the dependent variable. Variance inflation factors (VIF) were calculated with the car (v.3.1.3) R package, and covariates with a VIF score above two were excluded from subsequent analysis due to high multicollinearity among variables in the baseline data of SARS-CoV-2-positive cohorts. To identify statistically significant DMTWs for each COVID-19 severity group, the Benjamini-

Hochberg (BH) method was applied to adjust p-values for multiple testing. Tiled windows with BH-adjusted p-value < 0.05 were deemed statistically significant. A one-sided Fisher test was performed to determine the significance of the overlap between DMTW, DMGs, and enriched biological pathways across different COVID-19 severity groups, while Pearson correlation coefficients were computed to evaluate the correlation between different severity groups. Statistical significance was denoted using the following asterisk notations: *** for p < 0.001, ** for p < 0.01, * for p < 0.05, “.” for p < 0.1, and no symbol for p ≥ 0.1.

## Results

### Multicollinearity assessment and baseline data of SARS-CoV-2 positive cohorts

Variance inflation factor (VIF) analysis did not show significant multicollinearity among variables, except for relationships involving COVID-19 (Supplementary Figure S3). The clinical cohorts enrolled in the study are shown in Figure 1. The demographic characteristics of all participants (excluding samples that failed quality control) are summarized in Table 1.

**Table 1.**
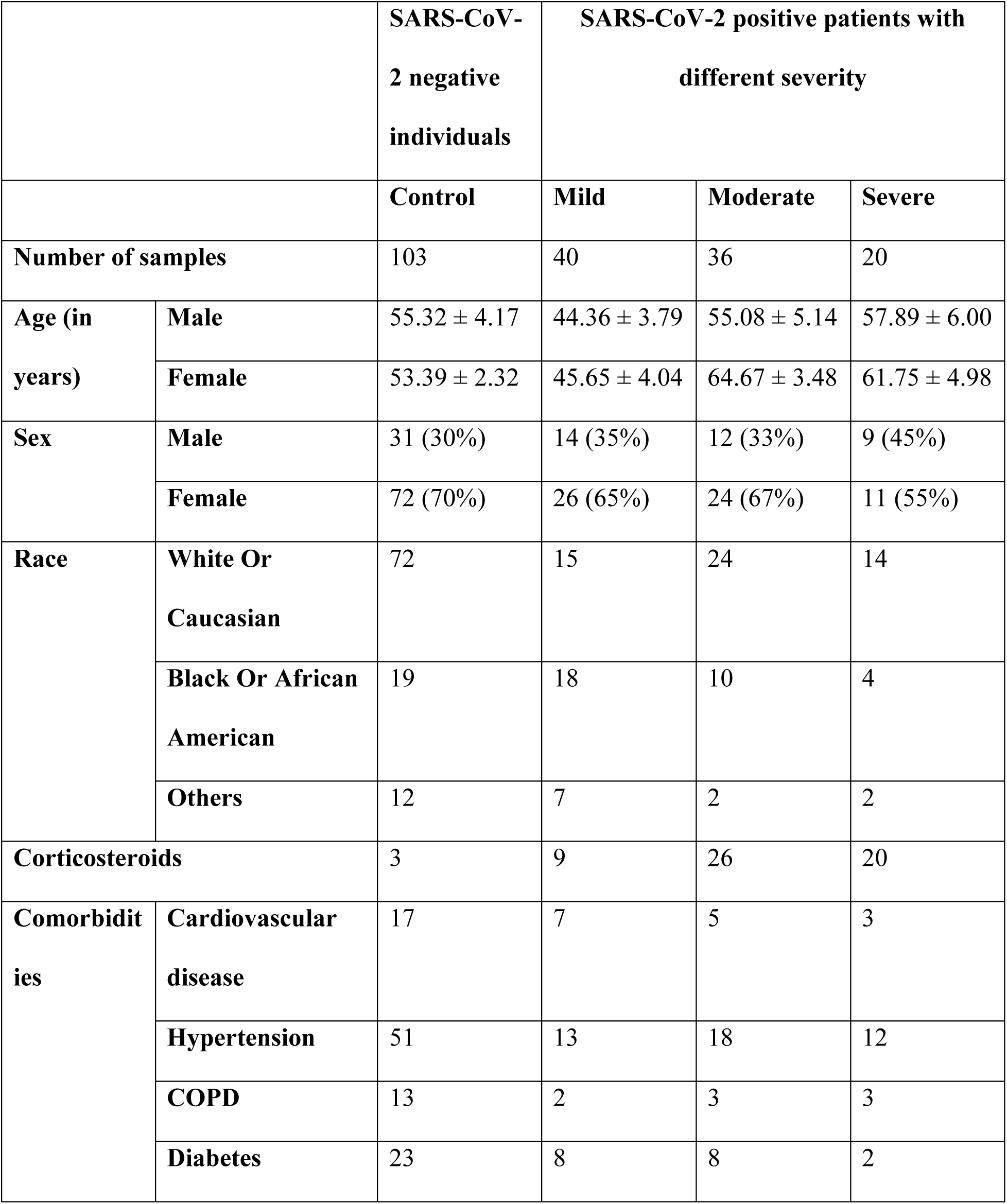
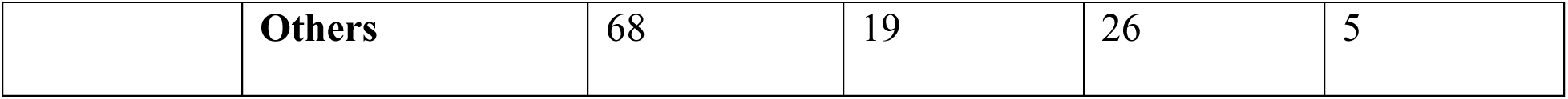
Demographic and baseline characteristics of SARS-CoV-2 positive (COVID-19) and SARS-CoV-2 negative (control) subjects. Demographic characteristics were calculated after excluding samples that did not pass whole-genome bisulfite sequencing (WGBS) quality control or were identified as outliers during data preprocessing. “Others” comorbidities include cancer, chronic kidney disease, depression, immunodeficiency, and obesity. Age is presented as mean ± SEM.

|  |  | **SARS-CoV-2 negative individuals** | **SARS-CoV-2 positive patients with different severity** |  |  |
| --- | --- | --- | --- | --- | --- |
|  |  | **Control** | **Mild** | **Moderate** | **Severe** |
| **Number of samples** |  | 103 | 40 | 36 | 20 |
| **Age (in years)** | **Male** | 55.32 ± 4.17 | 44.36 ± 3.79 | 55.08 ± 5.14 | 57.89 ± 6.00 |
|  | **Female** | 53.39 ± 2.32 | 45.65 ± 4.04 | 64.67 ± 3.48 | 61.75 ± 4.98 |
| **Sex** | **Male** | 31 (30%) | 14 (35%) | 12 (33%) | 9 (45%) |
|  | **Female** | 72 (70%) | 26 (65%) | 24 (67%) | 11 (55%) |
| **Race** | **White Or Caucasian** | 72 | 15 | 24 | 14 |
|  | **Black Or African American** | 19 | 18 | 10 | 4 |
|  | **Others** | 12 | 7 | 2 | 2 |
| **Corticosteroids** |  | 3 | 9 | 26 | 20 |
| **Comorbidities** | **Cardiovascular disease** | 17 | 7 | 5 | 3 |
|  | **Hypertension** | 51 | 13 | 18 | 12 |
|  | **COPD** | 13 | 2 | 3 | 3 |
|  | **Diabetes** | 23 | 8 | 8 | 2 |
|  | **Others** | 68 | 19 | 26 | 5 |

### Differential DNA methylation in the blood of SARS-CoV-2-positive patients reveals reduced methylation across the whole genome

We performed a genome-wide analysis to identify DNA methylation changes associated with SARS-CoV-2 infection. The Manhattan plot (Figure 2A-D) displays the distribution of differential methylation at individual genomic loci in all SARS-CoV-2-positive patients, followed by stratification based on their disease severity (mild, moderate, and severe) compared to the SARS-CoV-2-negative (control) group. To better understand the epigenetic landscape linked to SARS-CoV-2 infection, we analyzed DMTWs followed by tiling 4,843,203 windows using a linear DSS model. The analysis identified 78 hypomethylated and 6 hypermethylated windows in SARS-CoV-2-positive patients versus the controls. When stratified by disease severity, the number of hypomethylated windows was 25 in mild cases, 19 in moderate cases, and 42 in severe cases. Conversely, hypermethylated windows were limited, with 0, 4, and 5 identified in mild, moderate, and severe cases, respectively. These results suggest that hypomethylation increases with higher disease severity (Supplementary Table S1 A-D, Figure 3).

**Figure 2.**
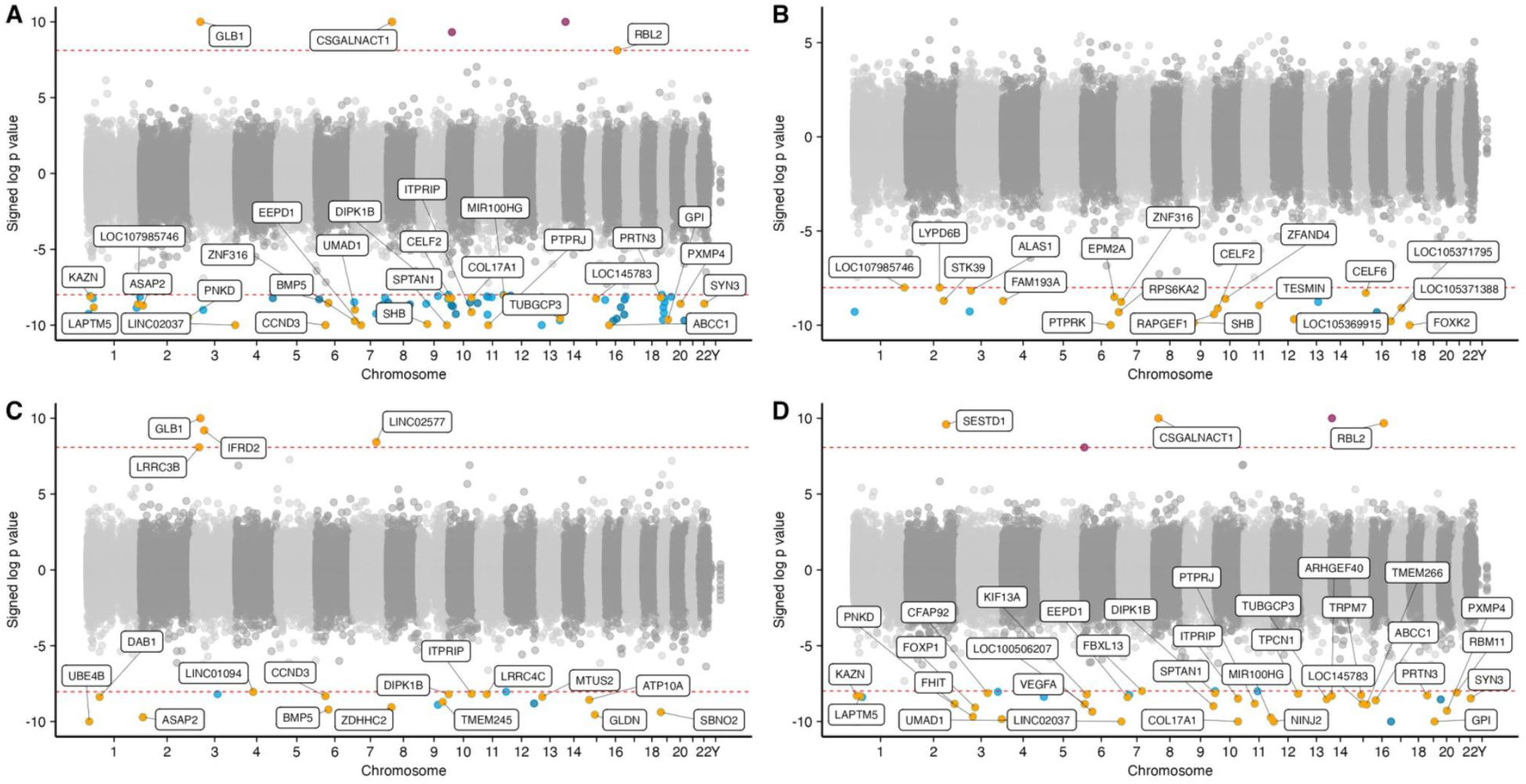
Manhattan plots showing differential DNA methylation across genomic loci in SARS-CoV-2-positive patients. (A) Comparison of all SARS-CoV-2-positive cohorts with SARS-CoV-2-negative controls. (B–D) Stratified analyses by COVID-19 disease severity: mild (B), moderate (C), and severe (D) cases compared to negative controls. The signed – log10 p-value reflects both statistical significance and direction of methylation change: positive values denote hypermethylation, while negative values indicate hypomethylation. Labeled points represent genes with significantly differentially methylated tiled windows (DMTWs) located within promoters, 1–5 kb upstream regions, or introns in each respective severity group.

**Figure 3.**
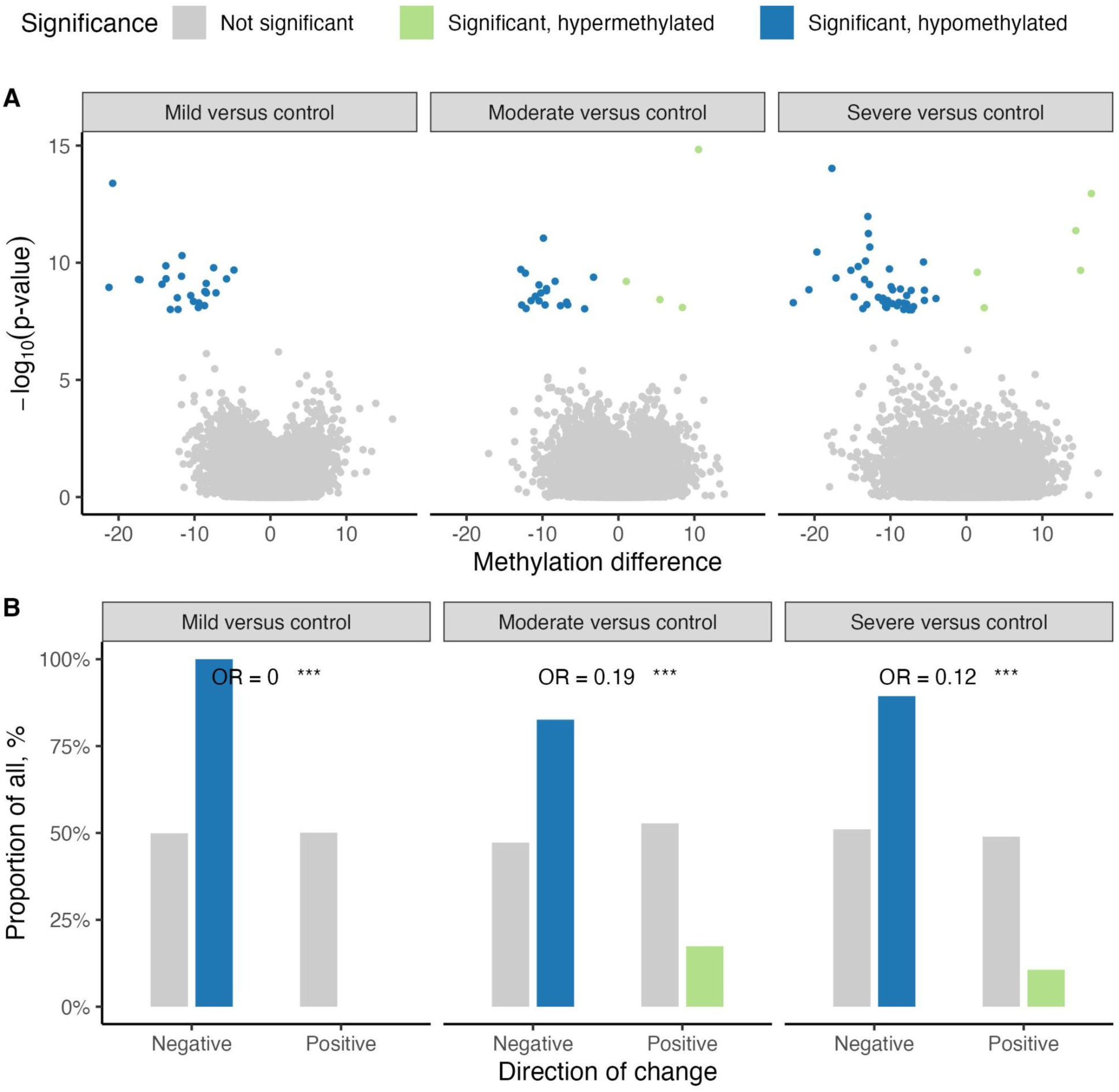
Differential DNA methylation in SARS-CoV-2-positive patients stratified by disease severity compared to negative controls. (A) Volcano plots illustrating the distribution of methylation differences in mild, moderate, and severe COVID-19 cases versus controls. Each point represents a methylation window; hypomethylated windows are shown in blue, no significant change in grey, and hypermethylated in green. (B) Bar plots illustrate the proportion of differentially methylated windows (DMTWs) (hypomethylated - blue and hypermethylated – green) in significant and non-significant (gray) groups. The odds ratio (OR) represents the result from Fisher’s exact test used to evaluate DMTWs. The asterisks denote statistical significance as *** for p < 0.001, ** for p < 0.01. Pathway enrichment analysis

### Identification of Differentially Methylated Genes (DMGs) in SARS-CoV-2-positive patients

DMTWs identified in SARS-CoV-2-positive patients with different disease severities versus controls were located within the promoters, 1-5 kb region upstream of the transcription start site (TSS), and intronic regions of various genes, which were categorized as DMGs. We found 19 DMGs in mild, 19 in moderate, and 35 in severe COVID-19 patients (FDR<0.05) (Table 2A). Notably, among these DMGs, 9, 8, and 15 in mild, moderate, and severe cases, respectively, are novel DMGs that have not been reported in previous epigenetic studies on COVID-19 (Table 2B). (4, 36).

**Table 2A.** Differentially Methylated Genes (DMGs) identified in SARS-CoV-2 positive patients. The table lists the DMGs in SARS-CoV-2 positive patients with mild, moderate, and severe cases, compared to negative controls. Gene locus indicates the genomic regions (e.g., promoter, intron, 1–5 kb upstream) with differential methylation, based on hg38 annotations. Methylation difference shows the Δβ (%) per group, where negative values indicate hypomethylation. FDR represents Benjamini– Hochberg–adjusted p-values.

| Symbol | Gene Locus | Methylation difference |  |  |  | FDR |  |  |  |
| --- | --- | --- | --- | --- | --- | --- | --- | --- | --- |
|  |  | COVID | Mild | Moderate | Severe | COVID | Mild | Moderate | Severe |
| ABCC1 | ABCC1<br>(hg38_genes_introns) /<br>ABCC1<br>(hg38_genes_promoters) | -5.51 | -4.22 | -4.44 | -7.86 | 0.0001 | 1.0000 | 0.4469 | 0.0122 |
| ALAS1 | ALAS1<br>(hg38_genes_1to5kb) /<br>ALAS1<br>(hg38_genes_exons) /<br>ALAS1<br>(hg38_genes_introns) | -8.67 | -8.67 | -9.59 |  | 0.4735 | 0.0327 | 1.0000 |  |
| ALAS1 | ALAS1<br>(hg38_genes_introns) /<br>ALAS1<br>(hg38_genes_1to5kb) |  |  |  | -8.93 |  |  |  | 1.0000 |
| ARHGEF40 | ARHGEF40<br>(hg38_genes_1to5kb) /<br>ARHGEF40<br>(hg38_genes_introns) | -5.74 | -<br>10.92 |  | -8.90 | 1.0000 | 1.0000 |  | 0.0239 |
| ARHGEF40 | ARHGEF40<br>(hg38_genes_promoters) /<br>ARHGEF40<br>(hg38_genes_exons) /<br>ARHGEF40<br>(hg38_genes_3UTRs) /<br>ARHGEF40<br>(hg38_genes_introns) |  |  | -2.22 |  |  |  | 1.0000 |  |
| ASAP2 | ASAP2<br>(hg38_genes_introns) | -11.06 | -9.13 | -12.85 | -8.46 | 0.0095 | 1.0000 | 0.0009 | 0.9750 |
| ATP10A | ATP10A<br>(hg38_genes_introns) /<br>ATP10A<br>(hg38_genes_1to5kb) |  | -6.98 |  |  |  | 1.0000 |  |  |
| ATP10A | ATP10A<br>(hg38_genes_introns) /<br>ATP10A<br>(hg38_genes_exons) /<br>ATP10A<br>(hg38_genes_5UTRs) | -6.59 |  | -10.93 | -7.35 | 0.1712 |  | 0.0131 | 1.0000 |
| BMP5 | BMP5<br>(hg38_genes_introns) | -4.87 | -5.28 | -8.34 | -4.01 | 0.0140 | 1.0000 | 0.0030 | 1.0000 |
| CCND3 | CCND3<br>(hg38_genes_introns) | -8.66 | -9.50 | -6.82 | -9.66 | 0.0003 | 1.0000 | 0.0232 | 1.0000 |
| CELF2 | CELF2<br>(hg38_genes_introns) |  |  | 5.59 |  |  |  | 1.0000 |  |
| CELF2 | CELF2<br>(hg38_genes_introns) /<br>CELF2<br>(hg38_genes_1to5kb) | -8.75 | -8.46 |  | -10.69 | 0.0288 | 0.0037 |  | 1.0000 |
| CELF6 | LOC105370887<br>(hg38_genes_introns) /<br>CELF6<br>(hg38_genes_introns) | -7.91 | -9.43 | -8.24 | -7.52 | 0.5898 | 0.0249 | 1.0000 | 1.0000 |
| CFAP92 | CFAP92<br>(hg38_genes_introns) | -7.43 | -5.03 |  | -10.63 | 1.0000 | 1.0000 |  | 0.0366 |
| CFAP92 | CFAP92<br>(hg38_genes_introns) /<br>CFAP92<br>(hg38_genes_exons) / |  |  | 4.72 |  |  |  | 1.0000 |  |
|  | CFAP92<br>(hg38_genes_promoters) |  |  |  |  |  |  |  |  |
| COL17A1 | COL17A1<br>(hg38_genes_exons) /<br>COL17A1<br>(hg38_genes_3UTRs) /<br>COL17A1<br>(hg38_genes_introns) | -9.07 |  | -8.69 | -13.29 | 0.0320 |  | 1.0000 | 0.0004 |
| COL17A1 | MIR936<br>(hg38_genes_1to5kb) /<br>COL17A1<br>(hg38_genes_introns) /<br>COL17A1<br>(hg38_genes_exons) |  | -9.66 |  |  |  | 1.0000 |  |  |
| CSGALNACT1 | CSGALNACT1<br>(hg38_genes_introns) | 10.23 | -6.66 | 8.37 | 16.45 | 0.0000 | 1.0000 | 0.0569 | 0.0000 |
| DAB1 | DAB1<br>(hg38_genes_exons) /<br>DAB1<br>(hg38_genes_3UTRs) |  | -<br>11.37 |  |  |  | 1.0000 |  |  |
| DAB1 | DAB1<br>(hg38_genes_introns) | -4.61 |  | -11.51 | -3.39 | 1.0000 |  | 0.0202 | 1.0000 |
| DIPK1B | DIPK1B<br>(hg38_genes_introns) | -5.75 | -5.85 | -6.67 | -6.96 | 0.0312 | 1.0000 | 0.0303 | 0.0362 |
| EEPD1 | EEPD1<br>(hg38_genes_introns) | -4.40 | -6.03 | -2.95 | -5.53 | 0.0003 | 0.5095 | 1.0000 | 0.0196 |
| EPM2A | EPM2A<br>(hg38_genes_introns) | -9.79 | -<br>12.27 | -12.69 | -10.51 | 1.0000 | 0.0152 | 1.0000 | 1.0000 |
| FAM193A | FAM193A<br>(hg38_genes_introns) | -4.43 |  | -10.51 | -6.85 | 1.0000 |  | 1.0000 | 1.0000 |
| FAM193A | FAM193A<br>(hg38_genes_introns) / |  | -7.17 |  |  |  | 0.0093 |  |  |
|  | FAM193A<br>(hg38_genes_1to5kb) |  |  |  |  |  |  |  |  |
| FBXL13 | FBXL13<br>(hg38_genes_introns) | -4.70 | -2.72 | -13.47 | -7.55 | 1.0000 | 1.0000 | 1.0000 | 0.0497 |
| FHIT | FHIT<br>(hg38_genes_introns) | -8.67 | 5.90 | -7.34 | -15.21 | 0.4237 | 1.0000 | 1.0000 | 0.0010 |
| FOXK2 | FOXK2<br>(hg38_genes_introns) | -7.75 | -11.65 | 5.78 | -11.47 | 1.0000 | 0.0002 | 1.0000 | 1.0000 |
| FOXP1 | FOXP1<br>(hg38_genes_introns) | -8.44 |  |  | -12.75 | 1.0000 |  |  | 0.0041 |
| FOXP1 | FOXP1<br>(hg38_genes_introns) /<br>EIF4E3<br>(hg38_genes_introns) |  |  | 8.90 |  |  |  | 1.0000 |  |
| FOXP1 | FOXP1<br>(hg38_genes_introns) /<br>FOXP1<br>(hg38_genes_1to5kb) |  | -6.49 |  |  |  | 1.0000 |  |  |
| GLB1 | GLB1<br>(hg38_genes_introns) | 7.08 | 5.49 | 10.55 | 5.20 | 0.0000 | 1.0000 | 0.0000 | 1.0000 |
| GLDN | GLDN<br>(hg38_genes_introns) | -4.36 | 4.55 | -12.25 | -7.92 | 1.0000 | 1.0000 | 0.0014 | 1.0000 |
| GPI | GPI<br>(hg38_genes_1to5kb) |  | -4.73 |  |  |  | 1.0000 |  |  |
| GPI | GPI<br>(hg38_genes_introns) | -7.93 |  | -4.13 | -12.91 | 0.0011 |  | 1.0000 | 0.0000 |
| IFRD2 | IFRD2<br>(hg38_genes_1to5kb) /<br>IFRD2<br>(hg38_genes_introns) /<br>IFRD2<br>(hg38_genes_exons) / | 0.62 | 0.84 | 1.04 | -0.02 | 0.7458 | 1.0000 | 0.0030 | 1.0000 |
|  | IFRD2<br>(hg38_genes_promoters) |  |  |  |  |  |  |  |  |
| ITPRIP | ITPRIP<br>(hg38_genes_1to5kb) | -7.16 |  | -7.63 | -10.98 | 0.0035 |  | 0.0329 | 0.0159 |
| ITPRIP | ITPRIP<br>(hg38_genes_introns) |  | -7.39 |  |  |  | 1.0000 |  |  |
| KAZN | KAZN<br>(hg38_genes_introns) | -6.15 | -3.77 | -7.03 | -22.80 | 0.0371 | 1.0000 | 0.1056 | 0.0247 |
| KIF13A | KIF13A<br>(hg38_genes_introns) |  | -5.56 | -5.74 |  |  | 1.0000 | 1.0000 |  |
| KIF13A | KIF13A<br>(hg38_genes_introns) /<br>KIF13A<br>(hg38_genes_exons) /<br>KIF13A<br>(hg38_genes_3UTRs) | -6.86 |  |  | -13.12 | 1.0000 |  |  | 0.0299 |
| LAPTM5 | LAPTM5<br>(hg38_genes_introns) /<br>MIR4420<br>(hg38_genes_1to5kb) | -7.32 | -4.05 | -6.92 | -10.82 | 0.0072 | 1.0000 | 1.0000 | 0.0227 |
| LINC01094 | LINC01094<br>(hg38_genes_introns) /<br>LINC01094<br>(hg38_genes_promoters)<br>/ LINC01094<br>(hg38_genes_exons) | -10.19 | -10.03 | -12.16 | -8.38 | 0.3261 | 1.0000 | 0.0437 | 1.0000 |
| LINC02037 | LINC02036<br>(hg38_genes_introns) /<br>LINC02037<br>(hg38_genes_promoters) | -9.68 | -8.36 | -6.44 | -14.25 | 0.0002 | 1.0000 | 1.0000 | 0.0007 |
| LINC02577 | LINC02577<br>(hg38_genes_introns) | 4.15 | 3.54 | 5.47 | 3.65 | 0.3288 | 1.0000 | 0.0181 | 1.0000 |
| LOC100506207 | LOC100506207<br>(hg38_genes_introns) |  | -7.72 |  |  |  | 1.0000 |  |  |
| LOC100506207 | LOC100506207<br>(hg38_genes_introns) /<br>LOC105374914<br>(hg38_genes_introns) /<br>LOC105374913<br>(hg38_genes_introns) /<br>LOC105374910<br>(hg38_genes_introns) | -10.75 |  |  | -20.74 | 0.2990 |  |  | 0.0069 |
| LOC100506207 | LOC100506207<br>(hg38_genes_introns) /<br>LOC112267952<br>(hg38_genes_introns) /<br>LOC105374914<br>(hg38_genes_introns) |  |  | 4.25 |  |  |  | 1.0000 |  |
| LOC105369915 | LOC105369915<br>(hg38_genes_introns) | -3.93 | -4.81 | 4.29 | -7.03 | 1.0000 | 0.0010 | 1.0000 | 1.0000 |
| LOC105371388 | LOC105371388<br>(hg38_genes_introns) /<br>LOC105371388<br>(hg38_genes_1to5kb) | -6.01 | -7.48 | -3.71 | -4.63 | 0.6497 | 0.0008 | 1.0000 | 1.0000 |
| LOC105371795 | LOC105371795<br>(hg38_genes_1to5kb) | -11.46 | -<br>14.28 | -8.67 | -11.42 | 1.0000 | 0.0040 | 1.0000 | 1.0000 |
| LOC107985746 | LOC107985746<br>(hg38_genes_introns) | -12.06 | -<br>13.18 | -9.82 | -13.18 | 0.0118 | 0.0478 | 1.0000 | 1.0000 |
| LOC145783 | ZNF280D<br>(hg38_genes_introns) /<br>LOC145783<br>(hg38_genes_introns) /<br>LOC105370833<br>(hg38_genes_promoters) | -3.38 | -2.95 | -1.65 | -5.53 | 0.0273 | 1.0000 | 1.0000 | 0.0072 |
|  | / LOC145783<br>(hg38_genes_exons) |  |  |  |  |  |  |  |  |
| LRRC3B | LRRC3B<br>(hg38_genes_1to5kb) /<br>LRRC3B-AS1<br>(hg38_genes_introns) /<br>LRRC3B-AS1<br>(hg38_genes_exons) |  |  |  | 10.36 |  |  |  | 1.0000 |
| LRRC3B | LRRC3B<br>(hg38_genes_introns) | 2.80 | -6.89 | 8.42 |  | 1.0000 | 1.0000 | 0.0393 |  |
| LRRC4C | LRRC4C<br>(hg38_genes_introns) | -6.12 | -6.59 | -9.66 | -5.68 | 1.0000 | 1.0000 | 0.0307 | 1.0000 |
| LYPD6B | LYPD6B<br>(hg38_genes_introns) /<br>LOC105373678<br>(hg38_genes_promoters) | -4.57 | -<br>12.17 | -3.50 |  | 1.0000 | 0.0478 | 1.0000 |  |
| LYPD6B | LYPD6B<br>(hg38_genes_introns) /<br>LYPD6B<br>(hg38_genes_1to5kb) |  |  |  | -3.51 |  |  |  | 1.0000 |
| MIR100HG | MIR100HG<br>(hg38_genes_introns) | -6.00 |  | -6.79 | -10.14 | 0.0434 |  | 1.0000 | 0.0009 |
| MIR100HG | MIR100HG<br>(hg38_genes_introns) /<br>MIR100HG<br>(hg38_genes_exons) |  | -5.59 |  |  |  | 1.0000 |  |  |
| MTUS2 | MTUS2<br>(hg38_genes_introns) | -5.18 | -6.11 | -10.45 | 9.10 | 1.0000 | 1.0000 | 0.0206 | 1.0000 |
| NINJ2 | NINJ2<br>(hg38_genes_introns) /<br>NINJ2<br>(hg38_genes_1to5kb) |  | 4.18 |  |  |  | 1.0000 |  |  |
| NINJ2 | NINJ2<br>(hg38_genes_introns) /<br>NINJ2-AS1<br>(hg38_genes_introns) /<br>NINJ2-AS1<br>(hg38_genes_exons) | -6.89 |  | -4.69 | -12.72 | 0.0908 |  | 1.0000 | 0.0001 |
| PNKD | PNKD<br>(hg38_genes_introns) /<br>CATIP-AS2<br>(hg38_genes_introns) /<br>PNKD<br>(hg38_genes_1to5kb) | -6.96 |  | -5.29 | -9.72 | 0.0016 |  | 1.0000 | 0.0070 |
| PNKD | PNKD<br>(hg38_genes_introns) /<br>PNKD<br>(hg38_genes_promoters) |  | -1.93 |  |  |  | 1.0000 |  |  |
| PRTN3 | ELANE<br>(hg38_genes_1to5kb) /<br>PRTN3<br>(hg38_genes_exons) /<br>PRTN3<br>(hg38_genes_3UTRs) | -5.51 |  | -6.33 | -8.24 | 0.0331 |  | 1.0000 | 0.0255 |
| PRTN3 | PRTN3<br>(hg38_genes_1to5kb) |  | -3.55 |  |  |  | 1.0000 |  |  |
| PTPRJ | PTPRJ<br>(hg38_genes_introns) | -4.79 | -6.66 | -4.11 | -7.28 | 0.0004 | 1.0000 | 1.0000 | 0.0073 |
| PTPRK | PTPRK<br>(hg38_genes_1to5kb) | -9.21 | -<br>20.77 |  |  | 1.0000 | 0.0000 |  |  |
| PTPRK | PTPRK<br>(hg38_genes_1to5kb) /<br>PTPRK-AS1<br>(hg38_genes_introns) /<br>PTPRK (hg38_genes_promoters) / PTPRK<br>(hg38_genes_introns) |  |  |  | -8.12 |  |  |  | 1.0000 |
| PTPRK | PTPRK<br>(hg38_genes_introns) / PTPRK<br>(hg38_genes_1to5kb) |  |  | 4.88 |  |  |  | 1.0000 |  |
| PXMP4 | PXMP4<br>(hg38_genes_1to5kb) | -9.08 |  | -7.74 | -13.41 | 0.0125 |  | 1.0000 | 0.0025 |
| PXMP4 | PXMP4<br>(hg38_genes_promoters) |  | -3.56 |  |  |  | 1.0000 |  |  |
| RAPGEF1 | RAPGEF1<br>(hg38_genes_exons) / RAPGEF1<br>(hg38_genes_introns) | -5.36 |  |  | -8.48 | 0.4121 |  |  | 0.2982 |
| RAPGEF1 | RAPGEF1<br>(hg38_genes_introns) |  | -11.71 | -7.18 |  |  | 0.0018 | 1.0000 |  |
| RBL2 | RBL2<br>(hg38_genes_introns) | 7.71 | 5.38 | 2.72 | 15.04 | 0.0369 | 1.0000 | 1.0000 | 0.0010 |
| RBM11 | RBM11<br>(hg38_genes_introns) / RBM11<br>(hg38_genes_1to5kb) | -5.75 |  | -5.51 | -10.51 | 1.0000 |  | 1.0000 | 0.0394 |
| RBM11 | RBM11<br>(hg38_genes_introns) / RBM11<br>(hg38_genes_promoters) |  | -5.90 |  |  |  | 1.0000 |  |  |
| RPS6KA2 | RPS6KA2<br>(hg38_genes_introns) |  | -5.78 | 6.89 |  |  | 0.0024 | 1.0000 |  |
| RPS6KA2 | RPS6KA2<br>(hg38_genes_introns) / RPS6KA2<br>(hg38_genes_1to5kb) / | -5.81 |  |  | -8.93 | 1.0000 |  |  | 1.0000 |
|  | RPS6KA2-IT1<br>(hg38_genes_introns) |  |  |  |  |  |  |  |  |
| SBNO2 | SBNO2<br>(hg38_genes_promoters)<br>/ SBNO2<br>(hg38_genes_introns) |  | -8.26 |  |  |  | 1.0000 |  |  |
| SBNO2 STK11 | STK11<br>(hg38_genes_1to5kb) /<br>SBNO2<br>(hg38_genes_1to5kb) /<br>SBNO2<br>(hg38_genes_introns) |  |  | -3.27 |  |  |  | 0.0020 |  |
| SBNO2 | STK11<br>(hg38_genes_promoters)<br>/ SBNO2<br>(hg38_genes_1to5kb) /<br>STK11<br>(hg38_genes_5UTRs) /<br>STK11<br>(hg38_genes_exons) | -7.03 |  |  | -8.75 | 1.0000 |  |  | 1.0000 |
| SESTD1 | SESTD1<br>(hg38_genes_introns) | -8.26 | -6.14 | -10.54 | 1.43 | 1.0000 | 1.0000 | 1.0000 | 0.0012 |
| SHB | SHB<br>(hg38_genes_introns) | -13.19 | -13.77 | -0.82 | -16.39 | 0.0006 | 0.0007 | 1.0000 | 0.5405 |
| SPTAN1 | SPTAN1<br>(hg38_genes_introns) | -10.09 | -8.75 | -11.60 | -9.91 | 0.0000 | 1.0000 | 0.1240 | 0.0051 |
| STK39 | STK39<br>(hg38_genes_introns) | -7.91 | -8.43 | -3.91 | -9.25 | 1.0000 | 0.0094 | 1.0000 | 1.0000 |
| SYN3 | SYN3<br>(hg38_genes_introns) | -2.92 | 10.48 | 9.57 | -4.01 | 0.0126 | 1.0000 | 0.1921 | 0.0163 |
| TESMIN | TESMIN<br>(hg38_genes_introns) | -8.12 |  | -8.79 | -8.93 | 1.0000 |  | 1.0000 | 1.0000 |
| TESMIN | TESMIN<br>(hg38_genes_introns) /<br>TESMIN<br>(hg38_genes_1to5kb) /<br>TESMIN<br>(hg38_genes_exons) |  | -<br>21.26 |  |  |  | 0.0054 |  |  |
| TMEM245 | TMEM245<br>(hg38_genes_1to5kb) | -8.53 |  | -10.21 |  | 0.1756 |  | 0.0095 |  |
| TMEM245 | TMEM245<br>(hg38_genes_introns) |  | 5.23 |  | -14.26 |  | 1.0000 |  | 1.0000 |
| TMEM266 | TMEM266<br>(hg38_genes_introns) | -5.49 | -4.90 | -2.81 | -8.69 | 0.6813 | 1.0000 | 1.0000 | 0.0064 |
| TPCN1 | TPCN1<br>(hg38_genes_introns) | -5.44 | -7.42 | -5.95 | -9.13 | 0.4750 | 1.0000 | 1.0000 | 0.0331 |
| TRPM7 | TRPM7<br>(hg38_genes_introns) | -6.76 | -6.69 | -5.95 | -9.77 | 0.1242 | 1.0000 | 1.0000 | 0.0270 |
| TUBGCP3 | TUBGCP3<br>(hg38_genes_introns) | -8.60 | 5.34 | -5.76 | -11.59 | 0.0013 | 1.0000 | 1.0000 | 0.0145 |
| UBE4B | UBE4B<br>(hg38_genes_introns) | -7.33 | -8.84 | -9.88 | -10.80 | 0.2153 | 1.0000 | 0.0000 | 1.0000 |
| UMAD1 | UMAD1<br>(hg38_genes_introns) |  | 3.95 |  |  |  | 1.0000 |  |  |
| UMAD1 | UMAD1<br>(hg38_genes_introns) /<br>RPA3<br>(hg38_genes_introns) | -8.92 |  | -8.36 | -12.98 | 0.0050 |  | 1.0000 | 0.0000 |
| VEGFA | VEGFA<br>(hg38_genes_1to5kb) /<br>VEGFA<br>(hg38_genes_exons) /<br>VEGFA<br>(hg38_genes_introns) |  | -5.21 |  |  |  | 1.0000 |  |  |
| VEGFA | VEGFA<br>(hg38_genes_1to5kb) /<br>VEGFA<br>(hg38_genes_introns) /<br>VEGFA<br>(hg38_genes_promoters) | -10.80 |  | -8.45 | -17.18 | 0.3259 |  | 1.0000 | 0.0022 |
| ZDHHC2 | ZDHHC2<br>(hg38_genes_introns) | -5.12 |  | -10.46 |  | 1.0000 |  | 0.0043 |  |
| ZDHHC2 | ZDHHC2<br>(hg38_genes_introns) /<br>ZDHHC2<br>(hg38_genes_1to5kb) |  | -6.17 |  |  |  | 1.0000 |  |  |
| ZDHHC2 | ZDHHC2<br>(hg38_genes_introns) /<br>ZDHHC2<br>(hg38_genes_5UTRs) /<br>ZDHHC2<br>(hg38_genes_exons) |  |  |  | -8.19 |  |  |  | 1.0000 |
| ZFAND4 | ZFAND4<br>(hg38_genes_1to5kb) /<br>ZFAND4<br>(hg38_genes_introns) |  |  | 7.95 |  |  |  | 1.0000 |  |
| ZFAND4 | ZFAND4<br>(hg38_genes_introns) /<br>ZFAND4<br>(hg38_genes_1to5kb) | -9.27 | -<br>10.50 |  |  | 1.0000 | 0.0122 |  |  |
| ZFAND4 | ZFAND4<br>(hg38_genes_promoters) |  |  |  | 1.87 |  |  |  | 1.0000 |
| ZNF316 | ZNF316<br>(hg38_genes_1to5kb) | -7.26 | -8.57 | -5.63 | -7.57 | 0.0009 | 0.0083 | 1.0000 | 1.0000 |

**Table 2B.** Novel Differentially Methylated Genes (DMGs) identified in SARS-CoV-2 positive patients. The table shows the novel DMGs identified in our cohort with mild, moderate, and severe COVID-19 compared to negative controls. These DMGs were found with differential methylation in different regions, including promoter, intron, and 1-5 kb upstream of the transcription start site. Methylation differences are expressed as the Δβ (%) per group, with negative values indicating hypomethylation. FDR represents Benjamini–Hochberg–adjusted p-values.

| Symbol | Gene Locus | Methylation difference | FDR |
| --- | --- | --- | --- |
| **Mild** |  |  |  |
| EPM2A | EPM2A (hg38_genes_introns) | -12.27 | 0.0152 |
| LOC105369915 | LOC105369915 (hg38_genes_introns) | -4.81 | 0.0010 |
| LOC105371388 | LOC105371388 (hg38_genes_introns) / LOC105371388 (hg38_genes_1to5kb) | -7.48 | 0.0008 |
| LOC105371795 | LOC105371795 (hg38_genes_1to5kb) | -14.28 | 0.0040 |
| LOC107985746 | LOC107985746 (hg38_genes_introns) | -13.18 | 0.0478 |
| LYPD6B | LYPD6B (hg38_genes_introns) / LOC105373678 (hg38_genes_promoters) | -12.17 | 0.0478 |
| TESMIN | TESMIN (hg38_genes_introns) / TESMIN (hg38_genes_1to5kb) / TESMIN (hg38_genes_exons) | -21.26 | 0.0054 |
| ZFAND4 | ZFAND4 (hg38_genes_introns) / ZFAND4 (hg38_genes_1to5kb) | -10.50 | 0.0122 |
| ZNF316 | ZNF316 (hg38_genes_1to5kb) | -8.57 | 0.0083 |
| **Moderate** |  |  |  |
| BMP5 | BMP5 (hg38_genes_introns) | -8.34 | 0.0030 |
| CCND3 | CCND3 (hg38_genes_introns) | -6.82 | 0.0232 |
| DIPK1B | DIPK1B (hg38_genes_introns) | -6.67 | 0.0303 |
| IFRD2 | IFRD2 (hg38_genes_1to5kb) / IFRD2 (hg38_genes_introns) / IFRD2 (hg38_genes_exons) / IFRD2 (hg38_genes_promoters) | 1.04 | 0.0030 |
| LINC02577 | LINC02577 (hg38_genes_introns) | 5.47 | 0.0181 |
| MTUS2 | MTUS2 (hg38_genes_introns) | -10.45 | 0.0206 |
| TMEM245 | TMEM245 (hg38_genes_1to5kb) | -10.21 | 0.0095 |
| UBE4B | UBE4B (hg38_genes_introns) | -9.88 | 0.0000 |
| **Severe** |  |  |  |
| COL17A1 | COL17A1 (hg38_genes_exons) / COL17A1 (hg38_genes_3UTRs) / COL17A1 (hg38_genes_introns) | -13.29 | 0.0004 |
| DIPK1B | DIPK1B (hg38_genes_introns) | -6.96 | 0.0362 |
| GPI | GPI (hg38_genes_introns) | -12.91 | 0.0000 |
| KIF13A | KIF13A (hg38_genes_introns) / KIF13A (hg38_genes_exons) / KIF13A (hg38_genes_3UTRs) | -13.12 | 0.0299 |
| LINC02037 | LINC02036 (hg38_genes_introns) / LINC02037 (hg38_genes_promoters) | -14.25 | 0.0007 |
| LOC100506207 | LOC100506207 (hg38_genes_introns) / LOC105374914 (hg38_genes_introns) / LOC105374913 (hg38_genes_introns) / LOC105374910 (hg38_genes_introns) | -20.74 | 0.0069 |
| MIR100HG | MIR100HG (hg38_genes_introns) | -10.14 | 0.0009 |
| RBM11 | RBM11 (hg38_genes_introns) / RBM11 (hg38_genes_1to5kb) | -10.51 | 0.0394 |
| SESTD1 | SESTD1 (hg38_genes_introns) | 1.43 | 0.0012 |
| SPTAN1 | SPTAN1 (hg38_genes_introns) | -9.91 | 0.0051 |
| TMEM266 | TMEM266 (hg38_genes_introns) | -8.69 | 0.0064 |
| TRPM7 | TRPM7 (hg38_genes_introns) | -9.77 | 0.0270 |
| TUBGCP3 | TUBGCP3 (hg38_genes_introns) | -11.59 | 0.0145 |
| UMAD1 | UMAD1 (hg38_genes_introns) / RPA3 (hg38_genes_introns) | -12.98 | 0.0000 |
| VEGFA | VEGFA (hg38_genes_1to5kb) / VEGFA (hg38_genes_introns) / VEGFA (hg38_genes_promoters) | -17.18 | 0.0022 |

Many DMGs, both new and previously reported, have been linked to a wide range of neurological conditions. Specifically, in mild cases: CELF2 (also known as CUGBP2), PTPRK, RAPGEF1, and STK39; in moderate cases: MTUS2, LRRC3B, and GLB1; and in severe cases: ARHGEF40, VEGF4, ABCC1, TPCN1, TUBGCP3, SYN3, and UMAD1 have been associated with risk for neurodegenerative disorders.

### Biological pathways influenced by differential DNA methylation in response to SARS-CoV-2 infection

Gene Set Enrichment Analysis (GSEA) revealed extensive epigenetic remodeling associated with SARS-CoV-2 infection, affecting over 400 biological pathways (Supplementary Table S2A-D). Most enriched pathways were driven by hypomethylated genes. In addition to the identified DMGs, differential methylation in SARS-CoV-2-positive patients significantly impacted pathways related to neurological processes, alongside the classical immune and inflammatory signaling.

Stratification based on severity further revealed a distinct pattern of pathway enrichment. In mild cases, hypomethylation-driven alterations were significantly enriched in pathways related to Alzheimer’s disease-amyloid secretase (PANTHER pathways database; NES = -2.057) as well as serotonin signaling and anxiety-related mechanisms (Wiki pathway database; NES = -2.228). In moderate cases, enrichment was observed in dopamine transport (GOBP database; NES = -2.279) and dopamine neurogenesis pathways (Wiki pathway database; NES = -2.759). Severe cases showed pronounced enrichment in pathways implicated in Parkinson’s disease (PANTHER pathway database; NES = -2.057). (Figure 4A-C, Supplementary Table S2A–D).

**Figure 4.**
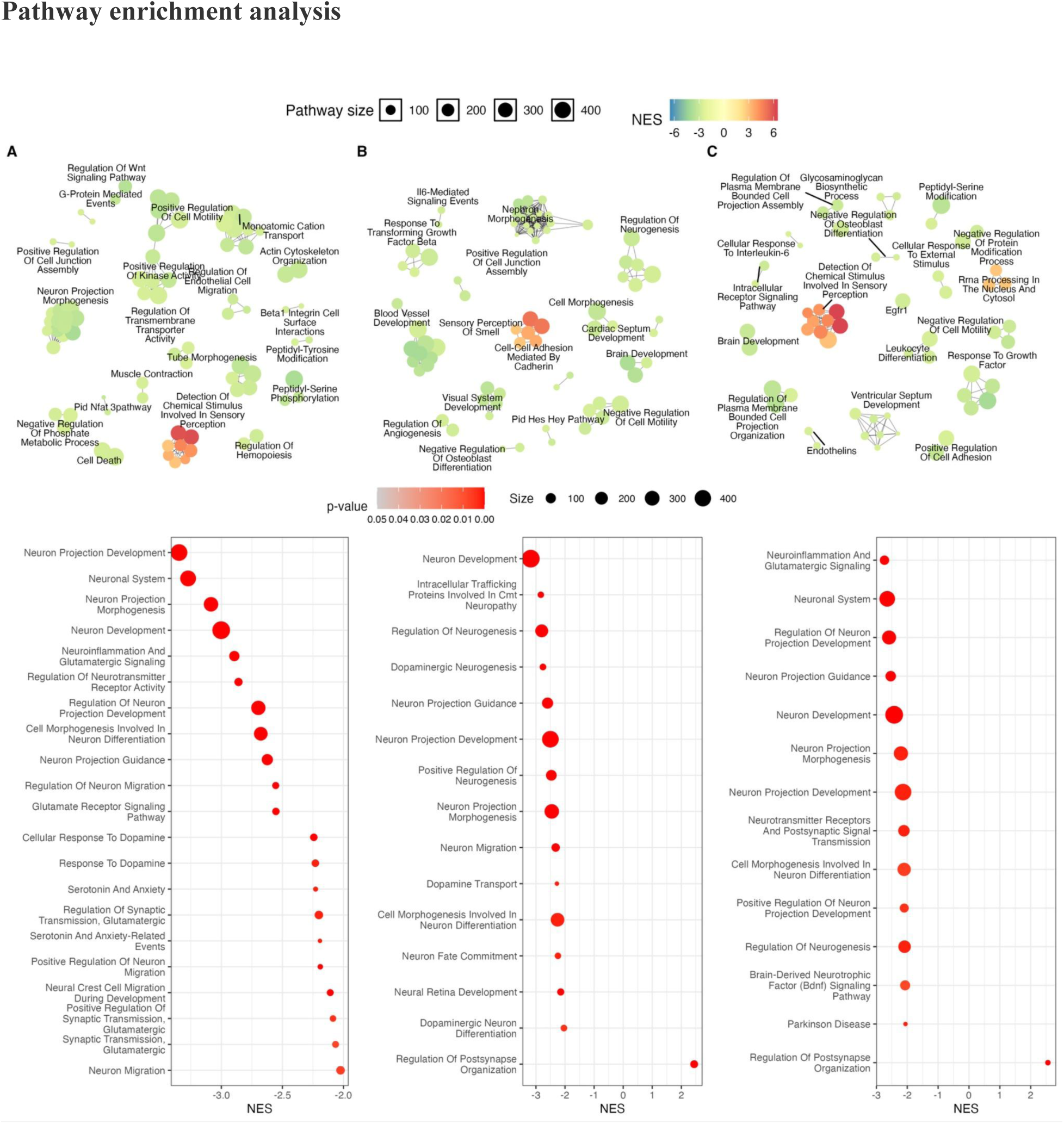
Gene Set Enrichment Analysis (GSEA) of pathways. The pathways altered by the differential methylation in SARS-CoV-2-positive patients with (A) mild, (B) moderate, and (C) severe symptoms are represented through an enrichment map (upper panel) and bubble plot (lower panel). The enrichment map displays a network and clustering of the top 100 significantly affected pathways identified through gene set enrichment analysis (GSEA). Each cluster is labeled with its most prominent pathway, as determined by the PageRank algorithm. Node color corresponds to the normalized enrichment score (NES), with negative values indicating a tendency for genes in the pathway to be hypomethylated. Node size reflects the number of genes identified within each pathway. The bubble plot highlights the most significantly enriched pathways related to neurodegenerative and psychiatric conditions, with bubble size proportional to gene set size and color indicating statistical significance (p < 0.05).

### Specificity of SARS-CoV-2 DNA methylation signature across mild, moderate, and severe cases

To identify specific epigenetic markers associated with severity in response to SARS-CoV-2 infection, we performed an intersection analysis of DMTWs and DMGs, which revealed minimal overlap across severity groups. Specifically, there was a 1.1% overlap between DMTWs (Figure 5A, Supplementary Table S3) and only 2.8% overlap in DMGs (Figure 5B, Table 2A) between moderate and severe cases. This 2.8% overlap in DMGs constitutes DIPK1B (FAM69B) and ITPRIP genes. Conversely, no overlap was observed in DMTW and DMGs between mild and severe or mild and moderate cases.

**Figure 5.**
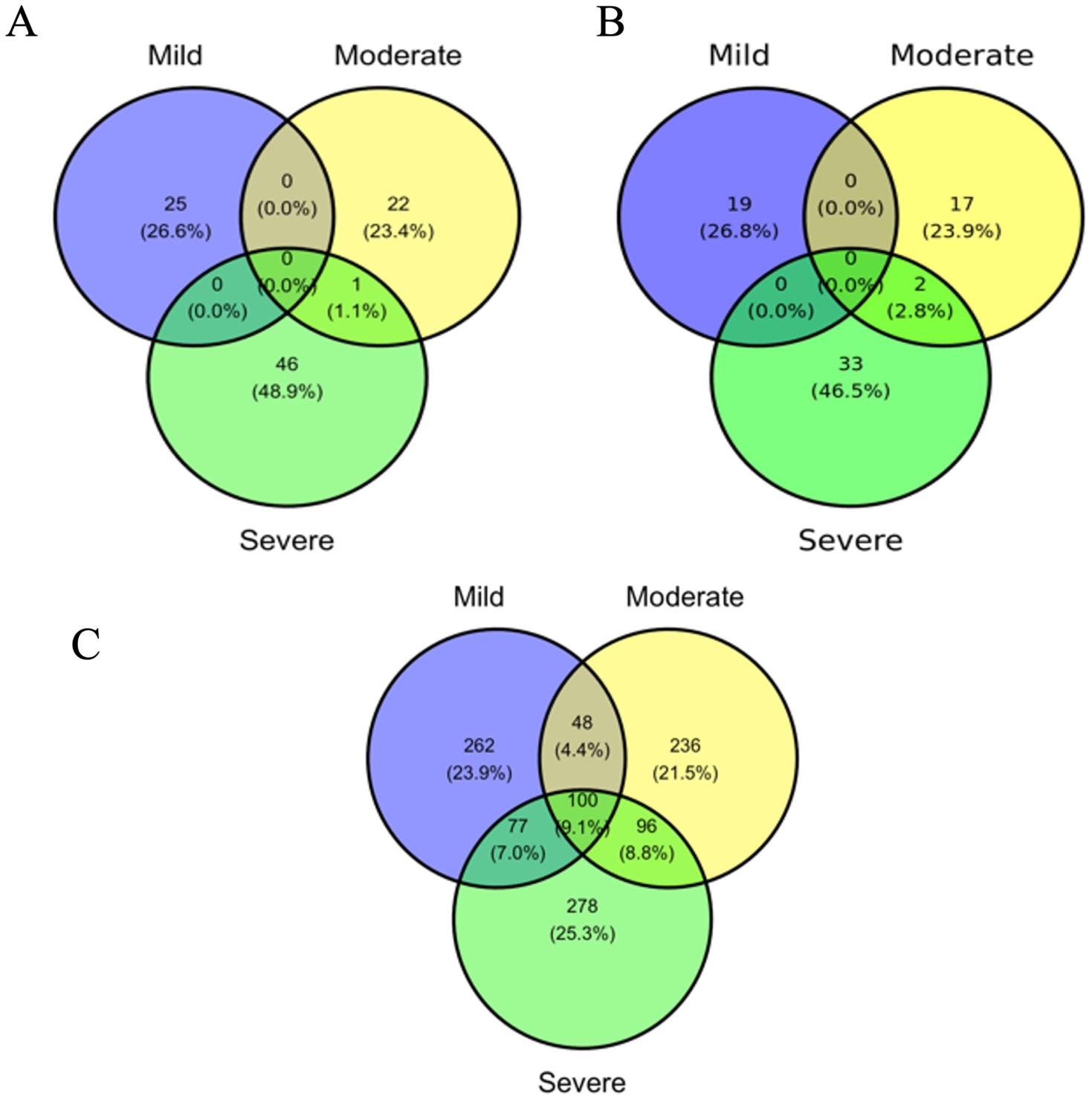
Intersection of DMTWs, DMGs, and pathways across the three COVID-19 severity groups. The Venn diagram illustrates the overlap among SARS-CoV-2-positive patients with mild, moderate, and severe symptoms for A) differentially methylated windows (DMTWs) and (B) differentially methylated genes (DMGs), and C) pathways affected by methylation changes. The diagrams highlight the shared and unique methylation signatures and pathway-level alterations across severity groups.

Further intersection analysis of significantly enriched biological pathways across mild, moderate, and severe cases revealed that most pathways were specific to each severity group, accounting for 23.9%, 21.5%, and 25.3%, respectively. Overlap between groups was limited, with 4.4%, 7.0%, and 8.8% of pathways shared between mild and moderate, mild and severe, and moderate and severe cases, respectively. Among the overlapping pathways, significant hypomethylation was observed in pathways related to axon and synapse assembly processes between mild and moderate cases. Similarly, pathways involved in neuroinflammation and glutamatergic signaling were enriched between mild and severe cases, while neurogenesis and postsynapse organization pathways were shared between moderate and severe cases. Only 9.1% of pathways were commonly enriched across all severity levels (Figure 5C, Supplementary Table S4). Interestingly, among those 9.1% of commonly enriched pathways, axon guidance and neuron projection-related processes, including guidance, development, and morphogenesis, were consistently hypomethylated, while pathways associated with olfactory processes, including olfactory receptor expression, translocation, and signaling, were hypermethylated across all severity groups.

### Correlation analysis of significantly DMTWs, gene order, and pathway enrichment in SARS-CoV-2-positive patients

Correlation analysis showed limited concordance in methylation patterns across COVID-19 severity groups. Based on signed log p-value (SLP), DMTWs exhibited low correlations of 0.26 for mild and moderate, 0.24 for mild and severe, and 0.30 for moderate and severe cases (Figure 6A). Similarly, the correlation between gene ranks was low, with coefficients of 0.082, 0.088, and 0.201 for the respective comparisons. (Figure 6B). In contrast, pathway enrichment correlations were moderate, with coefficients of 0.29, 0.30, and 0.41 for mild vs moderate, mild vs severe, and moderate vs severe cases, respectively. (Figure 6C).

**Figure 6.**
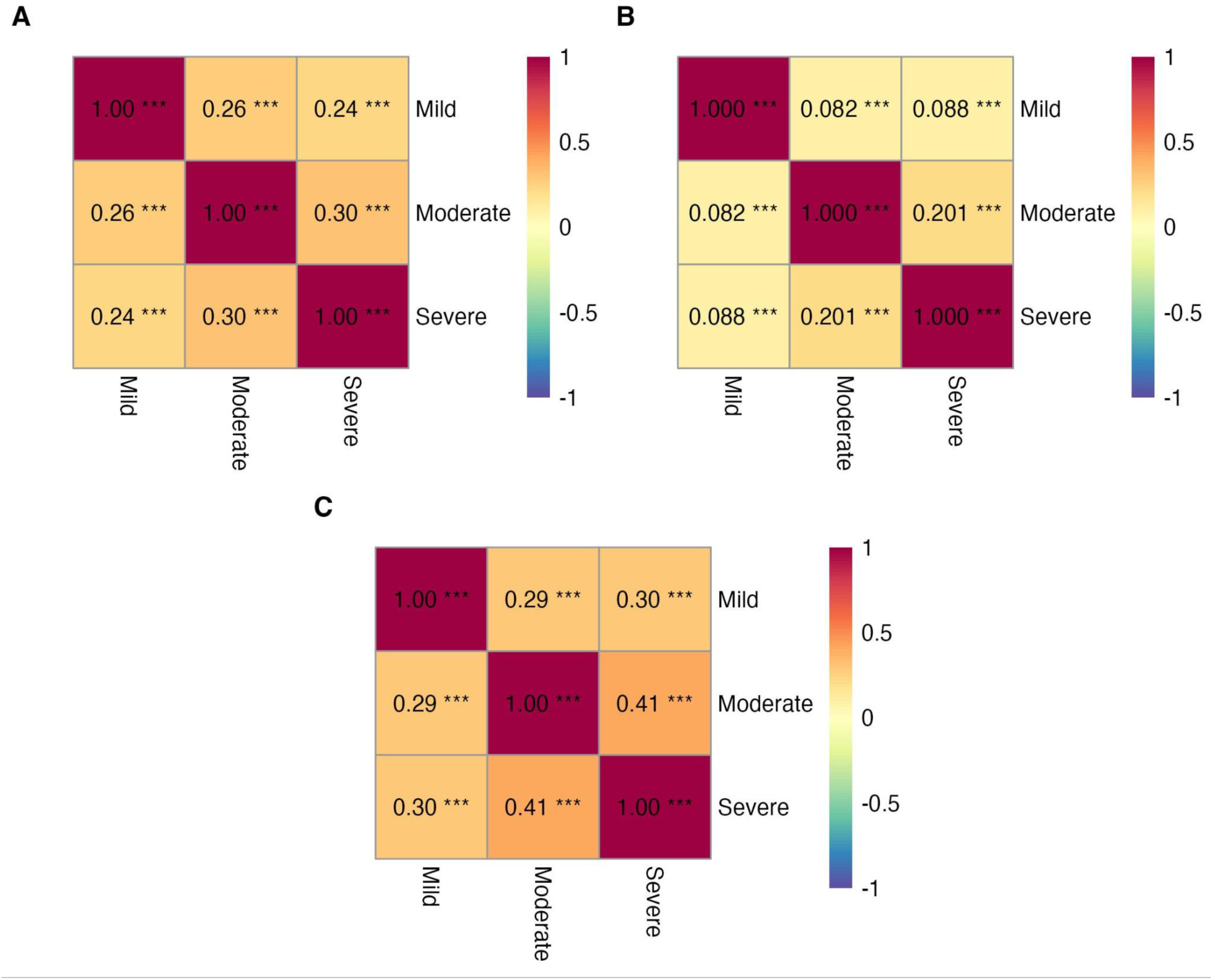
Correlation heatmap of differential methylation analysis in SARS-CoV-2 positive patients with varying disease severity. (A) Differentially methylated windows (DMTWs), (B) gene order, and (C) pathway enrichment in patients with mild, moderate, and severe symptoms. The heatmap shows minimal correlations, suggesting limited similarity in methylation changes across severity groups in response to SARS-CoV-2 infection. The color in the heatmap represents Pearson’s correlation coefficient. Statistical significance is denoted as follows: *** for p < 0.001, ** for p < 0.01, * for p < 0.05, . for p < 0.1, and no symbol for p ≥ 0.1.

## Discussion

We here utilized whole-genome bisulfite sequencing (WGBS) to characterize DNA methylation patterns in peripheral blood from SARS-CoV-2-positive patients compared to controls. Overall, we found a pronounced hypomethylation in the blood of the SARS-CoV-2-positive patients and identified 19 differentially methylated genes in patients with mild-, 19 in patients with moderate- and 35 differentially methylated genes in patients with severe symptoms. Multiple genes that were differentially methylated between SARS-CoV-2-positive patients and controls have been reported to be involved in neurodegenerative diseases.

WGBS provides a more comprehensive view of DNA methylation changes than array-based methods. It allows for the identification of novel differentially methylated regions not only in promoter regions but also in non-coding and intergenic regions, offering a broader perspective of the epigenomic landscape. Using WGBS on peripheral blood samples, we here examined DNA methylation changes in SARS-CoV-2-positive patients, stratified by disease severity, compared to SARS-CoV-2-negative controls. Our results demonstrate that SARS-CoV-2 infection is associated with significant epigenetic changes in peripheral blood. Compared to controls, we observed genome-wide hypomethylation in the blood samples of SARS-CoV-2-positive patients across different severities. These epigenetic modifications were enriched in genes and biological pathways related to neurodegenerative and psychiatric disorders, as well as immune, inflammatory, and viral responses.

Our findings enhance understanding of the wider effects of coronavirus infections. Decades before the COVID-19 pandemic, both SARS-CoV and Middle East Respiratory Syndrome (MERS-CoV) were shown to impact multiple organ systems, with documented neurological conditions such as encephalomyelitis and multiple sclerosis beyond their primary respiratory issues. (37–39). Both SARS-CoV and SARS-CoV-2 are part of the *Sarbecovirus* subgenus in the Coronaviridae family and are known to infect many different host species. (40, 41) and diverse cell types, including cells of the central nervous system CNS (42, 43). Reports of neurological symptoms such as seizures, extrapyramidal movement disorders, and mental health disorders in COVID-19 patients (44–46), have supported the hypothesis that SARS-CoV-2 can have long-term neurological effects. These effects may result from direct viral entry into the CNS through peripheral nerves, olfactory pathways, and the presence of angiotensin-converting enzyme (ACE2) receptors in glial and neural cells (47, 48). Postmortem studies in COVID-19 patients have shown the presence of the SARS-CoV-2 virus in the brain (49, 50), which per se could directly contribute to neurological disturbances. However, the neurological symptoms observed in COVID-19 may also stem from systemic inflammation or the synergistic effects of ischemia, coagulopathy, or hypoxia, even without any direct viral invasion of the brain (51, 52). Therefore, gaining a detailed understanding of the exact mechanisms behind the neurological effects in COVID-19 is lacking.

Several epigenetic studies have highlighted the role of DNA methylation in modulating immune responses, leukocyte activation, and host-pathogen interactions in COVID-19 patients (4, 18, 36, 53). In line with these studies, our WGBS analysis also observed similar changes in SARS-CoV-2-positive patients compared to negative controls. Additionally, our main goal was to identify the epigenetic modifications that may be linked to neurodegenerative and neuropsychiatric conditions in COVID-19 patients. Epigenetic changes in peripheral blood cells can reflect molecular alterations occurring in brain tissues (54, 55). Therefore, we specifically examined whole-genome DNA methylation in the blood samples of COVID-19 patients with varying disease severity to identify epigenetically altered genes and biological pathways involved in neurodegenerative and neuropsychiatric disorders. These findings may provide initial insights into potential peripheral epigenetic biomarkers for neurological involvement or disease prognosis.

The observed genome-wide hypomethylation trend in SARS-CoV-2-positive patients, which we observed was more pronounced in severe cases and aligned with broader evidence of immune and epigenetic dysregulation during viral infections. Hypomethylation within various regions of numerous genes (DMGs), especially in promoter regions and 1-5 kb upstream regulatory regions, is noteworthy because these regions often contain transcription factor binding sites and enhancer elements that regulate gene expression (56). Our findings support previous studies suggesting that severe viral infections can alter the epigenetic landscape of immune and inflammatory cells, as well as host defense (4, 36). However, a key feature of our findings is the enrichment of DMGs that also relate to neurodegenerative diseases, as corroborated by multiple lines of prior genetic, transcriptomic, and experimental evidence. This indicates a connection between SARS-CoV-2-induced immune-epigenetic changes and molecular pathways involved in brain health, which has not previously been observed in detail.

Specifically, a subset of DMGs with hypomethylation in promoters, 1-5 kb upstream regulatory regions, and intronic regions has previously been reported in association with neurological conditions like Alzheimer’s disease and vascular dementia. In mild cases, CELF2 (CUGBP Elav-like Family Member 2 or CUGBP2) and PTPRK (Protein Tyrosine Phosphatase Receptor Type K) were hypomethylated in regulatory regions. CELF2 is known to interact with the apolipoprotein E (APOE) gene in familial late-onset Alzheimer’s disease (57). In vitro studies further demonstrated that CELF2 influences TREM2 (Triggering Receptor Expressed on Myeloid cells 2)-mediated microglial response to amyloid-beta (58). Similarly, PTPRK is involved in T-cell development, synaptic signaling, and neuronal connectivity and has been suggested as a susceptibility gene for Alzheimer’s disease and multiple sclerosis (59, 60). In severe cases, genes including VEGFA (Vascular Endothelial Growth Factor A), ARHGEF40 (Rho Guanine Nucleotide Exchange Factor 40), and ABCC1 (ATP Binding Cassette Subfamily C member 1) showed hypomethylation within promoters and upstream regions. VEGFA and ARHGEF40 are involved in neurovascular remodeling and have been identified as risk genes for AD and vascular dementia (61–63). ABCC1, a crucial transporter involved in leukotriene export and inflammation, has been previously associated with amyloid clearance and AD progression (64–67).

Another cluster of DMGs, especially those with differential methylation in the intronic region, is known to be altered in Parkinson’s disease and synaptic dysfunctions. In mild cases, STK39 (Serine Threonine Kinase 39), a stress-activated kinase, was hypomethylated, which has been shown to worsen anxiety-like behavior and is a risk gene for Parkinson’s disease (68–70). RAPGEF1 (Rap Guanine Nucleotide Exchange Factor 1), also altered in mild cases, has been associated with cerebrospinal fluid (CSF) alpha-synuclein levels and is predicted to be a downstream target of miRNAs involved in amyloid signaling (71, 72). In moderate cases, MTUS2 (Microtubule-Associated Scaffold Protein 2), altered in Parkinson’s disease and late-onset Alzheimer’s disease, and GLB1 (Galactosidase Beta 1), a lysosomal enzyme linked to neurodegeneration and glial dysfunction, were hypomethylated in the intronic region (73–76). LRRC3B (leucine-rich repeat-containing 3B), also observed in moderate cases, was previously found to be downregulated in the substantia nigra of Parkinson’s disease patients, which is the brain region that is most severely affected in Parkinson’s disease (77). Simultaneously, in severe cases, we identified hypomethylation within the intronic regions of genes, including SYN3 (Synapsin 3), which is known as a modulator of dopamine release and alpha-synuclein aggregation, and TPCN1 (Two Pore Segment Channel 1), which is involved in calcium signaling and linked to Alzheimer’s disease (78, 79). Additionally, severe cases were identified with altered TUBGCP3 (Tubulin Gamma Complex Component 3) and UMAD1 (UBAP1-MVB12-Associated (UMA) Domain Containing 1) genes. which have been reported as risk loci for Parkinson’s disease and Alzheimer’s disease, respectively (80, 81).

While most of the focus in epigenetic studies often lies in promoters and upstream regulatory regions due to their critical role in regulating gene expression, our analysis also identified significant changes in the intronic region, which is traditionally considered as a non-coding region but increasingly recognized for regulatory functions. Studies suggest an inverse correlation between DNA-methylation and both mRNA intron retention and gene expression (82, 83). Genes such as *RAPGEF1, STK39, MTUS2, LRRC3B, GLB1, TPCN1, TUBGCP3, SYN3,* and *UMAD1,* discussed above, were among those with hypomethylation in intron regions. The convergence of promoters, upstream regulators, and intronic changes in genes linked to neurodegenerative diseases indicates that SARS-CoV-2 infection might disrupt multiple regulatory layers of gene expression, impacting both initiation and post-transcriptional processing of essential transcripts.

Interestingly, while many DMTWs and DMGs were unique to specific severity groups, only a small subset overlapped between moderate and severe cases. This distinct distribution highlights the importance of stratifying COVID-19 patients by disease severity, as each group seems to have unique epigenetic signatures. Although all groups likely share a core immune-epigenetic response to SARS-CoV-2, the differences in methylation profiles suggest different biological mechanisms may underlie mild, moderate, and severe COVID-19. Among the few overlapping DMGs between moderate and severe cases were DIPK1B (Divergent Protein Kinase Domain 1B), also known as FAM69B, and ITPRIP (Inositol 1,4,5-Trisphosphate Receptor Interacting Protein), genes previously associated with autism spectrum disorders and synaptic transmission, respectively (84) (85). These common changes in more advanced disease stages may indicate overlapping neurological or synaptic pathways relevant to long-term neurological effects in severe COVID-19

Following the identification of DMGs, the pathway enrichment analysis might offer insights into the functional effects of the widespread hypomethylation observed across the SARS-CoV-2 severity groups. Besides the expected involvement of immune, inflammatory, and viral responses, angiogenesis, and blood coagulation (4, 18, 36, 86), our results reveal a significant epigenetic impact on CNS-related pathways. Many of these neurological pathways were enriched with hypomethylated genes. These observations suggest that epigenetic changes might go beyond systemic inflammation and could potentially affect neurological processes either directly or indirectly.

In line with the patterns observed in DMGs, mild cases exhibited hypomethylation-driven disruption of the Alzheimer’s disease-amyloid secretase pathway, suggesting a possible link between early neurodegenerative changes and COVID-19. This aligns with a systematic review discussing overlapping pathologies between cognitive decline in COVID-19 patients and Alzheimer’s disease-related dementia (87). Additional hypomethylation of pathways related to serotonin signaling and anxiety further supports the growing evidence documenting the increasing prevalence of psychiatric symptoms, including anxiety and depression, in individuals recovering from COVID-19 (88, 89). Moderate cases exhibited epigenetic disruptions specifically in the dopamine-transport and neurogenesis pathways, which may be linked to reported movement disorders in COVID-19 patients (90, 91). These changes indicate that the dopaminergic system, known for being highly sensitive to inflammatory insults, may also be vulnerable to SARS-CoV-2-induced epigenetic dysregulations. In severe cases, pathway dysregulation extended to Parkinson’s disease, which aligns with reports suggesting Parkinsonism could be a potential post-COVID-19 sequel (92–96). Although the different biological pathways affected in mild, moderate, and severe cases they all point to a shared neurological theme involved in neurodegeneration and disruptions in neurotransmitter systems. This suggests a continuum of severity-related epigenetic remodeling at the functional pathway level. While most differentially methylated pathways were specific to COVID-19 severity, a small subset—particularly those involved in olfactory processes, immune and inflammatory regulation, cellular signaling, and endocytosis—was consistently altered across all severity groups. This consistency potentially indicates their central role in COVID-19 pathogenesis (97–103). Notably, persistent hypermethylation in pathways related to olfactory processes, including olfactory receptor expression, translocation, and signaling, which was observed across all groups, aligns with the high prevalence of anosmia in COVID-19 patients (104, 105). This change is especially important given new evidence that olfactory problems often come before motor and cognitive issues in neurodegenerative diseases like Alzheimer’s and Parkinson’s disease (106–109). Complementing these findings, our intersection analysis revealed overlapping neurological disruptions among all severity groups, including hypomethylation of axon guidance and neuron projection-related processes. These pathways are critical for neural circuit formation and plasticity and have been implicated in neurodegenerative and psychiatric disorders (110). In Alzheimer’s and Parkinson’s disease, altered axon guidance signaling contributes to synaptic loss and dopaminergic dysfunction, respectively (111, 112).

## Limitations

This study was conducted at one single timepoint when patients were either positive or negative for SARS-CoV-2, and as such, we cannot determine for certain whether the infection caused the different methylation patterns, or whether patients with different epigenetic patterns were vulnerable to develop COVID-19 of different severity even before the infection. It is also possible that the observed epigenetic differences were unrelated to the infection. Therefore, the implicated genes and mechanisms deserve further attention in future longitudinal cohort studies as well as in pre-clinical mechanistic studies. Another important limitation is the use of whole blood for our epigenome analysis, which cannot fully serve as a proxy for brain-related epigenetic changes. However, whole blood can be employed for this purpose, as has been done in several other epigenome-wide studies addressing CNS dysfunctions (54, 55, 113–115).

## Conclusion

Our study shows that SARS-CoV-2 infection is associated with widespread, severity-specific epigenetic changes in peripheral blood, with minimal overlap in differentially methylated regions and genes among mild, moderate, and severe cases. Although correlations between groups were significant, their low magnitude infers a biologically meaningful divergence, supporting the view of COVID-19 as a spectrum of distinct syndromes rather than a single disease entity. While all groups showed disruption in the classical immune and inflammatory responses, each displayed unique features and distinct epigenetic profiles. The enrichment of genes and pathways associated with Alzheimer’s disease, Parkinson’s disease, and neuropsychiatric conditions could indicate a mechanistic connection between SARS-CoV-2 infection and neurological effects through ongoing epigenetic remodeling. However, it is important to note that our study is not causative by nature and only indicates associations. Our findings emphasize the importance of severity-based stratification and support further longitudinal research to investigate potential long-term neurological and psychiatric sequelae of COVID-19.

## Additional file 1

**Supplementary Figure S1 - Distribution of imputed methylation data**

Visualization of beta and M values calculated to assess methylation differences among SARS-CoV-2 positive patients with varying disease severities following removal of outliers.

**Supplementary Figure S2 - PCA and MDS analysis of DNA methylation**

PCA and MDS plots showing clustering of SARS-CoV-2 positive patient samples by symptom severity.

**Supplementary Figure S3 - Variance inflation analysis**

Heatmap showing Pearson correlation co-efficient to assess multicollinearity among baseline covariates.

## Additional file 2

**Supplementary Table S1. - Significant DMTWs SARS-CoV-2 positive cohorts**

The table lists the DMTWs identified as significant in SARS-CoV-2 positive patients with mild, moderate, and severe symptoms based on WGBS analysis.

## Additional file 3

**Supplementary Table S2. - Pathway enrichment**

The table summarizes enriched biological pathways in SARS-CoV-2-positive patients with different disease severity, as determined through pathway enrichment analysis.

## Additional file 4

**Supplementary Table S3. - Overlapped DMTWs**

The table provides the DMTWs that were found to overlap among SARS-CoV-2 positive patients with different disease severity.

## Additional file 5

**Supplementary Table S4. - Overlapped pathways**

The table shows the biological pathways that were commonly enriched among SARS-CoV-2-positive patients with different disease severity.

## Declaration

### Ethics approval and consent to participate

This study was approved by the Institutional Review Boards at Corewell Health East William Beaumont University Hospital (formerly Beaumont Health), Royal Oak, MI, USA (approval no. 2020-269) and Van Andel Institute, Grand Rapids, MI, USA (approval no. NHS 21014).

### Consent for publication

This study does not include any individual participant data (such as images or any personal details) in any form.

### Availability of data and materials

All data generated and/or analyzed during this study are included in this publication and its supplementary information files.

## List of abbreviations

COPD: Chronic obstructive pulmonary disorder
COVID-19: Coronavirus disease-19
DMGs: Differentially methylated genes
DMTWs: Differentially methylated tiling windows
GO: Gene Ontology
GSEA: Gene set enrichment analysis
GWAS: Genome-wide association studies
HumanCyc: Encyclopedia of Human Genes and Metabolism
KEGG: Kyoto Encyclopedia of Gene and Genome
MDS: Multi-Dimensional Scaling
MSigDB: Molecular Signature Database
NCL_Nature: National Cancer Institute Nature Pathway
NES: Normalized enrichment score
PANTHER: Protein Analysis Through Evolutionary Relationships
PCA: Principal Component Analysis
SARS-CoV-2: Severe acute respiratory syndrome coronavirus 2
SD: Standard deviation
VIF: Variance inflation factor

## Acknowledgement

We are thankful to the Farmer Family Foundation for the funds to accomplish the study. We also thank Van Andel Institute, Genomic Core, Grand Rapids (RRID: SCR_022913) for the generation of whole genome sequence libraries.

## Author’s contributions

S.Z. wrote the original draft of the manuscript. L.B. conceived and supervised the project. M.M., M.G., I.G., and J.G. contributed to methodology and bioinformatics analysis. S. Z. contextualized the bioinformatics findings. L.B., M.L.E.G., A.P., and P.B. provided guidance and interpretation of data. N.S. and S.F.G. contributed to the collection of human samples, S.Z., E.A., S.K., Q.S., and J.A.S. edited the manuscript and provided constructive feedback. All authors contributed to reviewing the final version of the manuscript. All authors read and approved the final manuscript.

