## Supplementary Figures for "Whole-genome DNA methylation profiling in COVID-19 positive patients reveals alterations in pathways linked to neurological dysfunction"

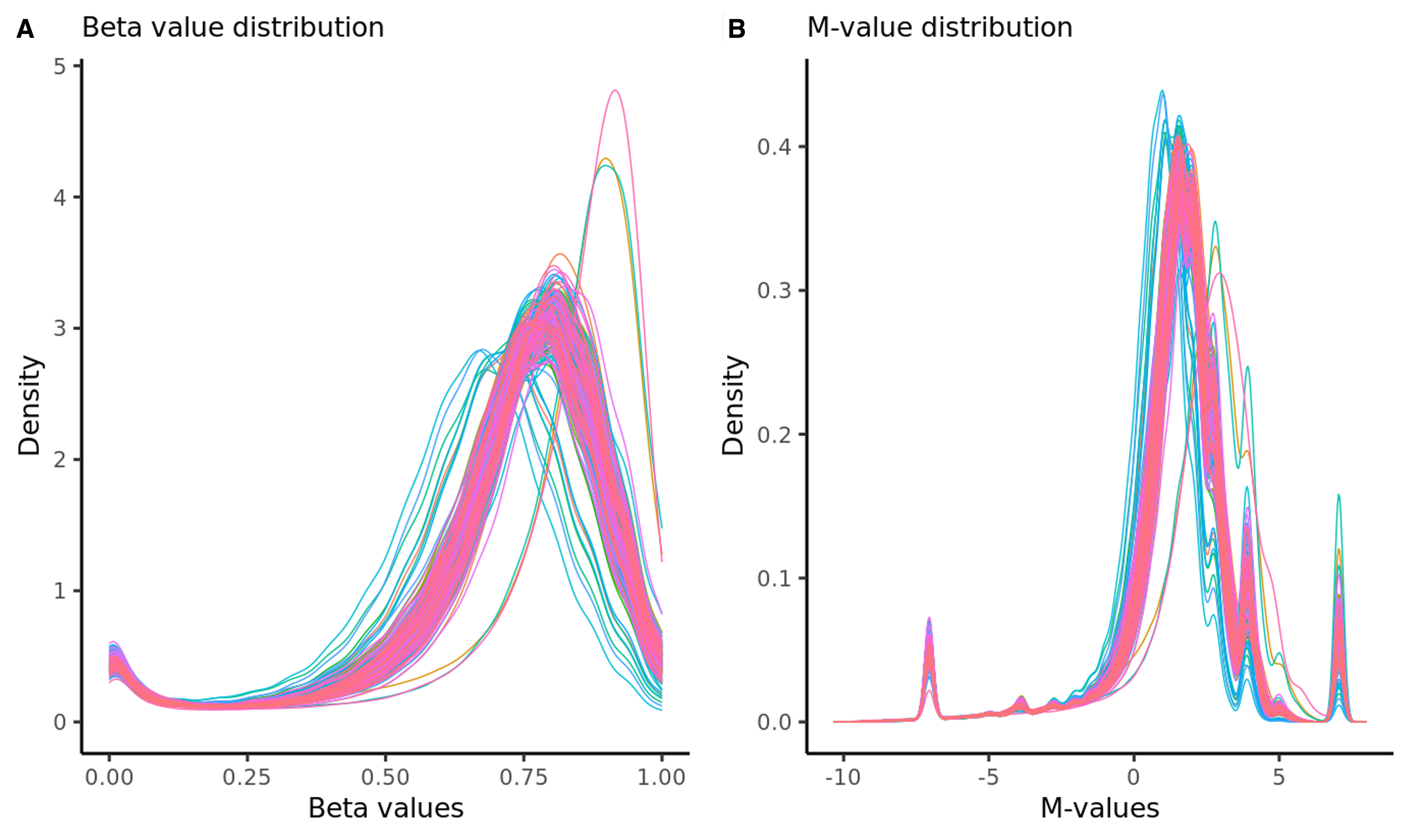

**Figure S1. Distribution of imputed data after outlier removal. (a) Beta and (b) M values were calculated to assess methylation differences among SARS-CoV-2 positive patients with varying levels of symptom severity.**

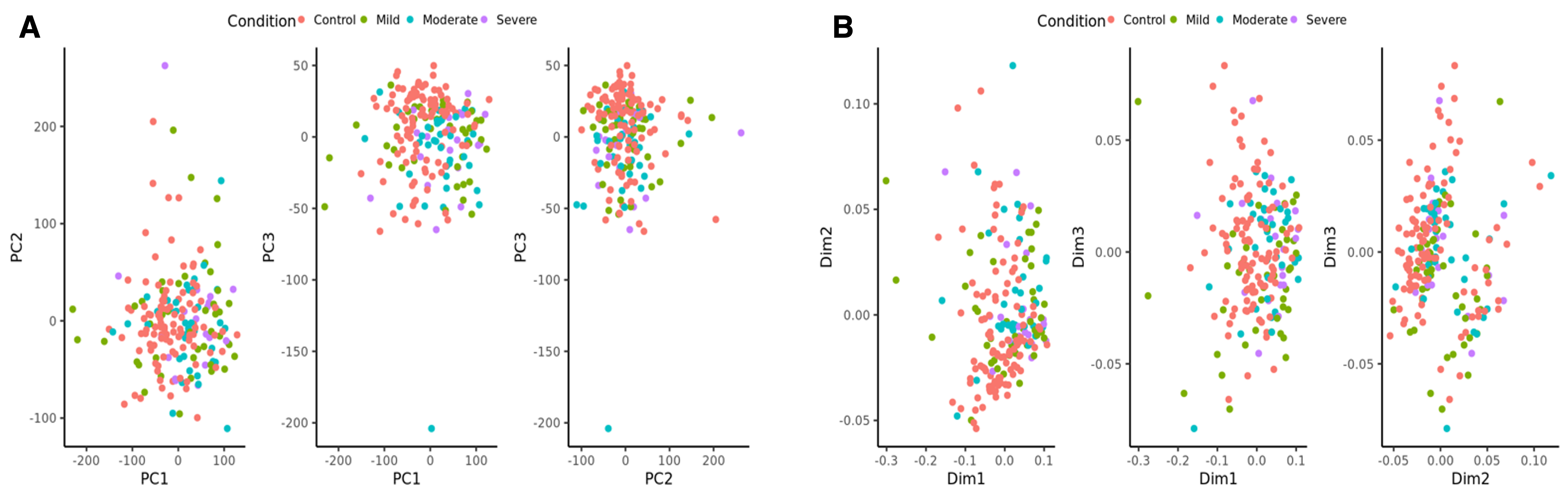

**Figure S2. Principal Component Analysis (PCA) and Multi-Dimensional Scaling (MDS) of DNA methylation data after outlier removal. A. The PCA plots depict the first three principal components (PC1, PC2, and PC3), showcasing the variance in the data across different SARS-CoV-2 symptom severity groups (mild, moderate, and severe). Each point represents a sample, with the grouping reflecting how the samples cluster based on their methylation profiles post-outlier removal. B. MDS plots illustrate the same samples, visualizing their clustering according to the underlying methylation profiles.**

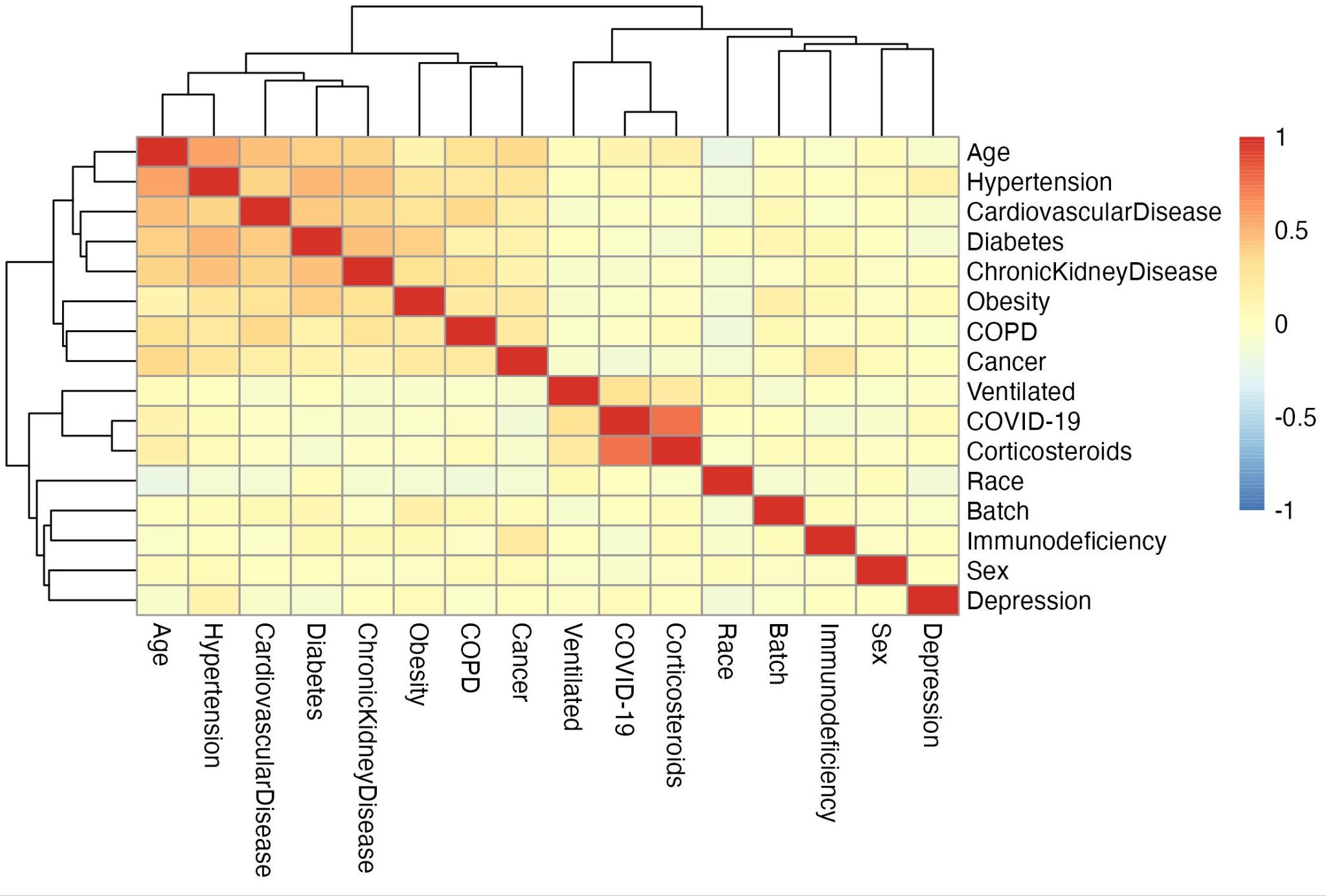

### Figure S3. Variance inflation function analysis to assess multicollinearity among variables in the baseline data of SARS-CoV-2 positive cohorts. Pairwise Pearson correlation coefficients were computed among all specified covariates. Cell color intensity reflects the magnitude and direction of the Pearson correlation coefficient, ranging from –1 (strong negative correlation) to +1 (strong positive correlation).
